# O_2_ photoreduction at acceptor side of Photosystem I provide photoprotection to conifer thylakoids in early spring

**DOI:** 10.1101/2022.10.21.513261

**Authors:** Pushan Bag, Tatyana Shutova, Dmitry Shevela, Jenna Lihavainen, Sanchali Nanda, Alexander G. Ivanov, Johannes Messinger, Stefan Jansson

## Abstract

Green organisms evolve O_2_ via photosynthesis and consume by respiration. Net O_2_ consumption only becomes dominant when photosynthesis is suppressed at night. Here, we show that green thylakoid membranes of Scots pine (*Pinus sylvestris L*) and Norway spruce (*Picea abies*) needles demonstrate strong O_2_ consumption even in the presence of light when extremely low temperatures coincide with high solar irradiation during early spring. This phenomenon deviates from the general finding that photosynthetic organisms evolve O_2_ upon illumination. By using different electron transport chain inhibitors, we showed that O_2_ consumption occurred around photosystem (PS) I and correlated with higher abundance of flavodiiron (Flv) A protein in ES thylakoid membranes. Furthermore, by measuring P700 absorption changes, we separated different alternative electron flow pathways and demonstrated that electron scavenging from the acceptor-side of PSI via O_2_ photoreduction is a major alternative pathway in ES. This photoprotection mechanism in vascular plants indicates that conifers have developed an adaptative evolution trajectory for growing in harsh environments.

## Introduction

The majority of O_2_ in the Earth’s atmosphere is generated by oceanic phytoplankton [1]–[3], which can achieve very high photosynthetic rates per weight biomass. O_2_ production/weight biomass of boreal conifer forests is low. However, their contribution to the global carbon balance and O_2_ production is still significant [4] since they are widespread in northern regions.

Photosynthetic O_2_ evolution from H_2_O splitting is carried out by a penta-μ-oxo bridged tetra-manganese calcium cluster (Mn_4_CaO_5_) in the oxygen-evolving complex (OEC) of photosystem (PS) II [5]–[7] during light reactions. The electron extracted from H_2_O is further transferred to PSI through several redox carriers and subsequently accepted by NADP^+^ to produce NADPH [8] in the photosynthetic electron transfer chain (PETC). Later, in the so-called dark reaction, CO_2_ assimilation occurs involving NADPH in the Calvin-Benson-Bassham (CBB) cycle [9]. The redox imbalance between light and dark reactions often leads to formation of reactive oxygen species (ROS), which can damage photosystems [10]. Hence, several protection mechanisms, such as non-photochemical quenching (NPQ) in PSII [11], [12], PTOX-mediated oxidation (chlororespiration) of the plastoquinone (PQ) pool [13], cyclic electron flow (CEF) [14], Mehler-like reactions (MR) [15], [16] around PSI and photorespiration (PR) via RuBisCO (ribulose bis-phosphate carboxylase oxygenase) in chloroplast stroma [17], have evolved in plants. PTOX, a chloroplastic non-heme diiron quinol oxidase, oxidizes the PQ pool via consumption of O_2_ [13], [18], whereas in CEF, a delta pH generated across thylakoid lumens enhances NPQ [19] and controls the excess electron flow towards PSI [20]. MR consume electrons from PSI to reduce O_2_ to H_2_O through H_2_O_2_ (hydrogen peroxide) [15], [21], thereby protecting PSI. In PR, RuBisCO fixes O_2_ instead of CO_2_ and releases CO_2_ through an inter-mitochondrial and inter-peroxisomal shuttle [17], [22]. Cyanobacteria [23]–[25], algae [26] and mosses [27] have an extra safeguard mechanism involving flavodiiron (Flv) proteins, known as pseudo CEF (PCEF), which can take up excess electrons from the PSI acceptor-side and reduce O_2_ directly to H_2_O.

In boreal forests, most plant species overwinter without exposing green parts to light, but conifers face extremely high oxidative stress in early spring (ES) when solar radiation is high but photosynthesis is constrained by low temperatures [10]. We recently demonstrated the molecular basis of the ‘sustained NPQ’ mechanism [28] that protects PSII in winter/ES in Scots pine [29] and Norway spruce [30]. Even though this quenching is extremely efficient in protecting PSII, it is hard to completely prevent light-driven ROS production under such low temperatures [31], [32]. Therefore, questions remain about the protection mechanism of PSI. Previous reports have suggested that PTOX [18], [33] and CEF may be involved [28], [34]. However, the absence of thylakoid NDH complexes (NADH dehydrogenase-like) [35], [36] in gymnosperms and the potential presence of an Flv-mediated PCEF [37] makes the situation complex. Moreover, PR has been shown not to be the major electron sink under low temperatures [38]. Obviously, conclusions made from studies of angiosperms may not hold true for conifers [39].

In the present study, we measured light induced O_2_ exchange in isolated thylakoids from summer (S) and ES in Scots pine and Norway spruce needles. Using different PETC inhibitors, such as DCMU (blocks Q_B_ site at PSII) [40] and mercuric chloride (HgCl_2_; profoundly affects electron transfer via plastocyanin) [21], [41], we obtained clear evidence that O_2_ photoreduction around PSI is much stronger than PSII O_2_ evolution in ES. In combination with P700 absorbance and immunodetection, our data suggest that in ES, O_2_ consumption via thylakoid bound Flv-dependent PCEF is a major functional electron sink. This mechanism could remove internal O_2_, prevent over-reduction of the acceptor side of PSI and may protect photosystems against photooxidative stress in ES better than other suggested pathways. Our study provides first functional evidence of this kind of photoreduction of O_2_ around PSI in vascular plants.

## Results

### Distinct dynamics of O_2_ evolution detected in thylakoid membranes of conifers and angiosperms

Isolated thylakoid membranes are devoid of stromal components, their illumination rapidly leads to over-reduction of the PQ pool, which lowers the rate of H_2_O splitting. Hence, to measure the “pure” O_2_-evolving activity of PSII without the influence of other components of the PETC [42], we performed O_2_ evolution measurements in the presence of PPBQ and FeCy, which accept electrons from the Q_B_ site on the PSII acceptor side (SI Appendix Fig. S1 and S2). First, we measured O_2_ exchange with a Clark type oxygen electrode (CTOE) (SI Appendix Fig. S1A, B). Second, we performed time resolved membrane inlet mass spectrometry (TR-MIMS) assays at 10% of H_2_^18^O enrichment (SI Appendix Fig. S2A, B) to discriminate between oxygen production (^16^O^18^O (*m/z* 34)) and consumption (mainly ^16^O_2_ (*m/z* 32)) reactions [42].

CTOE measurements showed that in spinach thylakoids, the O_2_ yield was independent of the O_2_ level in the medium as O_2_ exchange was similar in both air-saturated and O_2_-free buffer (Fig. 1A). In contrast, O_2_ evolution in S pine thylakoid membranes was strongly dependent on the O_2_ level in the medium (Fig. 1B, C, green line). O_2_ evolution of S pine thylakoids in air-saturated buffer was approximately half that of spinach thylakoids. Moreover, the O_2_ yield reached a maximum after the first 30 s (at 60^th^ second) and then decreased slowly over the next 30 s (60^th^ to 90^th^ second) of the illumination period (Fig. 1B, green line). In O_2_-free buffer, O_2_ evolution of S pine thylakoids increased for the first 50 s and then plateaued for the last 10 s (Fig. 1C, green line). Interestingly, spinach membranes in air-saturated buffer without PPBQ and FeCy supplementation did not show any O_2_ exchange (SI Appendix Fig. S1C), whereas S pine thylakoids showed only O_2_ consumption (SI Appendix Fig. S3, green line).

**Figure 1.**
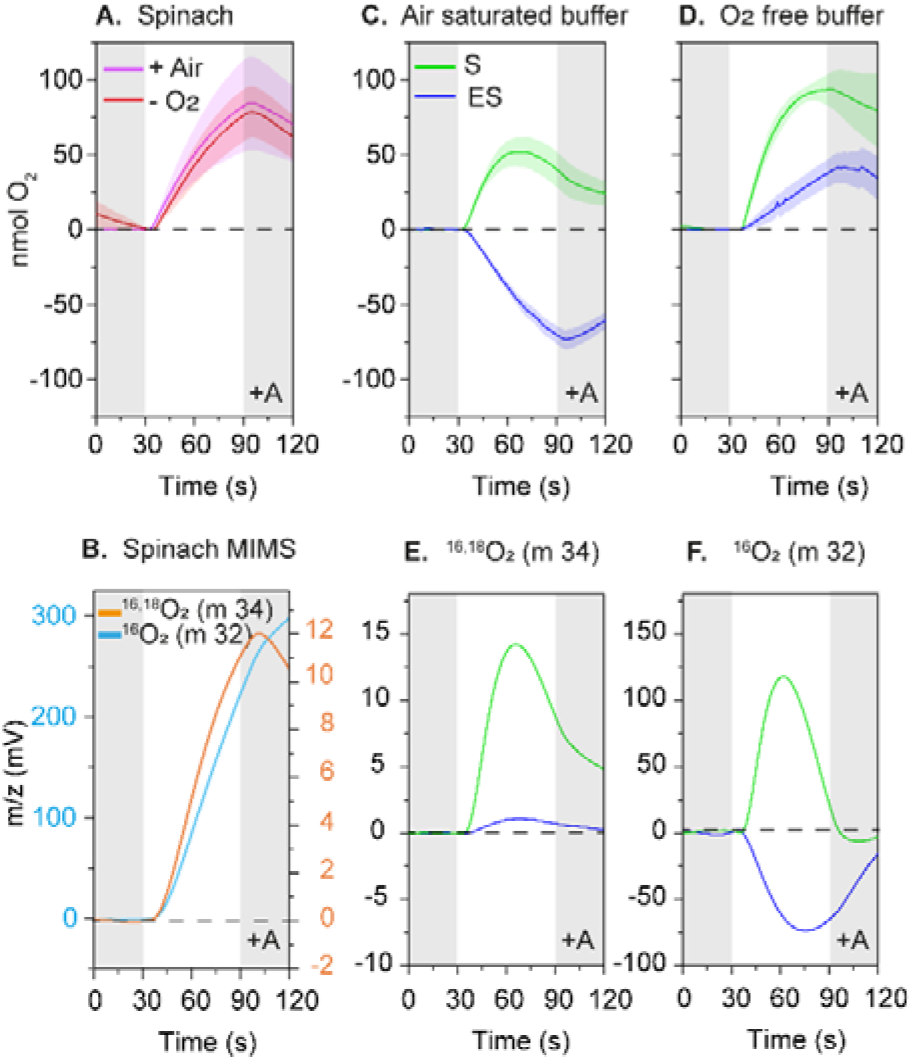
O_2_ yields in spinach and pine thylakoid membranes. **(A)** O_2_ yields in spinach thylakoid membranes in air-saturated and O_2_-free buffer measured with a CTOE. (**B)** ^16,18^O_2_ and ^16^O_2_ yields in spinach thylakoid membranes measured by TR-MIMS in partially degassed condition. **(C)** O_2_ yields in summer (S) and early spring (ES) pine thylakoid membranes in air, and (**D)** in O_2_-free buffer measured with a CTOE. **(E)** ^16,18^O_2_ and (**F)** ^16^O_2_ yields in S and ES pine thylakoid membranes under partially degassed conditions measured with TR-MIMS. Gray shaded regions indicate dark periods before and after illumination of the thylakoid membranes with 800 (CTOE) / 1200 (MIMS) μmol of photons m^−2^ s^−1^ for 60 seconds. In spinach, S and ES pine thylakoid membranes data represents O_2_ exchange corresponding to 50 μg of chlorophyll both in CTOE and MIMS measurements. For measurements in O_2_-free buffer, the thylakoid suspension buffer was bubbled with a continuous flow of N_2_ in the Clark electrode chamber until the O_2_ yield became zero prior to addition of the thylakoid membranes (for the experimental design, see SI Appendix Fig. S1 and S2;). Colored shaded regions around the average O_2_ yield curve indicate the standard deviation (*n*=3-4) in CTOE measurements. For TR-MIMS (10% H_2_ ^18^O enrichment), one representative spectrum out of 3-5 independent measurements is shown. Since the H_2_^18^O enrichment was small in TR-MIMS, the ^18,18^O_2_ trace was not considered for analysis. In all measurements, thylakoid membranes were supplemented with exogenous electron acceptors (+A, i.e., PPBQ and FeCy).

TR-MIMS demonstrated that both oxygen isotopolouges (^16^O_2_ and ^16,18^O_2_) exhibited similar O_2_ production kinetics (Fig. 1D) in spinach thylakoids. The distinct dynamics of O_2_ evolution in pine S thylakoids (as in CTOE measurements) were consistent with ^16^O_2_ and ^16,18^O_2_ yields in the MIMS experiment (Fig. 1E, F, green line). The results indicated that spinach and S pine thylakoid membranes had a similar maximum H_2_O oxidation capacity, but a possible O_2_ consuming process in S pine thylakoids became dominant over O_2_ evolution after approximately 30 s of illumination when O_2_ was available in the suspension.

### O_2_ consumption was dominant in early spring pine thylakoids

Photosynthetic activity has been shown to differ significantly in S and ES pine needles [29]. Therefore, we compared the light-dependent O_2_ exchange of ES and S pine thylakoid membranes using CTOE measurements. ES samples in air-saturated buffer showed strong O_2_ consumption (Fig. 1C, blue line). In O_2_-free buffer, O_2_ evolution in ES very slowly increased until the end of the illumination period, but the overall amplitude of ES samples was ~50% (Fig. 1D, blue line) of that of S samples (Fig. 1D, green line). This suggests that although light-dependent O_2_ evolution was present in ES, light-dependent O_2_ consumption was much stronger from the beginning, resulting in net consumption of O_2_. ES samples in air-saturated buffer without PPBQ and FeCy supplementation showed three times higher consumption than those from S (SI Appendix Fig. S3, blue line).

To understand the dynamics of simultaneous O_2_ evolution and consumption from a physiological perspective, we performed TR-MIMS on S and ES samples, which showed that ^16,18^O_2_ evolution in ES was only 10% of that in S (Fig. 1E). However, unlike in S, ^16^O_2_ was strongly consumed immediately after illumination and consumption reached a maximum after 40 s of illumination in ES (Fig. 1F). PSII showed typical ^16,18^O_2_ evolution immediately after illumination in both ES and S but with a much smaller amplitude in ES. This is consistent with ^16^O_2_ consumption due to processes occurring at later stages of the PETC.

### H_2_O oxidation was not the main source of electrons for photoreduction of O_2_ in ES thylakoids

All reduction reactions in the PETC require electrons, which are typically supplied by H_2_O oxidation in PSII. To determine whether electrons involved in the photo-consumption of O_2_ were supplied from PSII, we added DCMU to the thylakoid suspension. In presence of DCMU, the ^16,18^O_2_ yield diminished completely, suggesting that H_2_O oxidation in PSII was completely blocked in both S and ES (Fig. 2A). Surprisingly, both S and ES samples consumed ^16^O_2_, but the negative amplitude of ^16^O_2_ in ES was ~1.6 times to that of S (Fig. 2B). This shows that H_2_O oxidation in PSII did not supply electrons for the photoreduction of O_2_ neither in S nor ES. Hence, we concluded that in pine thylakoid membranes, simultaneous O_2_ evolution and consumption were two independent processes occurring in both S and ES but with different magnitudes. This difference resulted in contrasting net O_2_ yield patterns between S and ES.

**Figure 2.**
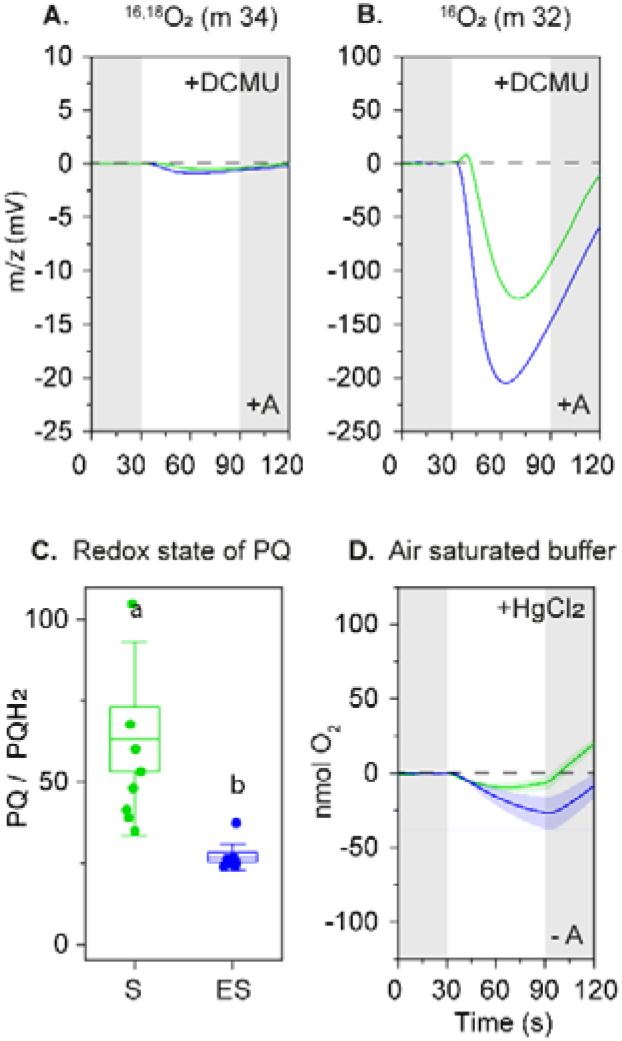
O_2_ yields in summer (S) and early spring (ES) pine thylakoid membranes in the presence of DCMU and redox state of the plastoquinone (PQ) pool. **(A)** ^16,18^O_2_ (note that O_2_ consumption in the mixed labeled O_2_ was small due to the low natural abundance of ^16,18^O_2_ compared to ^16^O_2_). (**B)** ^16^O_2_ yields in S and ES pine thylakoid membranes supplemented with 100 μM DCMU measured by TR-MIMS under partially degassed conditions in the presence of ‘A’ (‘+A’). Gray shaded regions indicate dark periods before and after illumination of the thylakoid membranes with 800 μmol of photons m^−2^ s^−1^ for 60 seconds. For TR-MIMS (10% H_2_^18^O enrichment), one representative spectrum out of 3-5 independent measurements is shown. **(C)** Redox state of the PQ pool presented as the ratio of plastoquinol (PQH2) to plastoquinone (PQ) measured using UPLC-APCI(−)QTOF-MS (*n*=9) (SI Appendix Table S2) from native state (dark adapted) S and ES thylakoid membranes. The whisker box indicates the SE and error bars indicate the standard deviation. Statistically significant mean differences were calculated by two-way ANNOVA (SI Appendix Table S5). **(D)** O_2_ yields in S and ES thylakoid membranes in air-saturated buffer measured by a CTOE with 1 mg ml^−1^ HgCl_2_ supplementation in the absence of ‘A’ (‘-A’). Note that the HgCl_2_ measurements were performed in absence of ‘A’ because ‘A’ takes electrons from Q_B_, which is upstream of plastocyanin - the major site of HgCl_2_ action - in the PETC. The green/blue colored shaded regions around the O_2_ yield curve indicates ±SE (*n*=3) in the CTOE measurements. In S and ES pine thylakoid membranes data represents O_2_ exchange corresponding to 50 μg of chlorophyll in both the CTOE and MIMS measurements. (For the experimental design, see SI Appendix Fig. S1 and S2,).

Earlier studies have suggested that high oxidative stress imposed on conifer needles in ES could lead to photoinhibition of PSII via photodamage [34], [43], [44]. Several mechanisms of photodamage have been suggested, and all involve a strong reduction in O_2_ evolution [45] (SI Appendix Fig S4 and the discussion there in). It has also been suggested that photodamage may form Mn-depleted PSII reaction centers that can consume O_2_ [40]. Our previous study of ES pine needles suggested substantial photoinhibition in PSII [29], but it was far from severe under the conditions tested. In the present study, ES thylakoids evolved ~50% O_2_ compared to S thylakoids (Fig. 1C). Hence, PSII was not severely photodamaged. This rules out the possibility that photodamaged PSII centres could be responsible for the O_2_ consumption in ES, and is in agreement with an O_2_ reduction site(s) downstream of PSII in the PETC.

### A reduced PQ pool supplied electrons to PSI for the photoreduction of O_2_

PQ, a common biological redox mediator, is present predominantly in the lumen of chloroplasts, mitochondria and the lipophilic compartment of bilayer membranes [46]. In higher plants, ~30% of the total PQ pool (fraction of the pool in the lipophilic compartment of bilayer thylakoid membranes) is known to be photochemically active [47]. If the electrons for O_2_ consumption in isolated ES thylakoid sample were supplied by the reduced PQ pool, it would be reflected in then redox state of the PQ pool present in the lipophilic compartment of the thylakoid bilayer. In the native state (dark adapted) photochemical PQ in ES thylakoids was in a predominantly reduced state: the PQ/PQH_2_ ratio in ES was ~37% lower than in S (Fig. 2C). In addition, the ubiquinone pool (UQ) was heavily reduced (SI Appendix Fig. S5). This suggests that the reduced PQ pool in ES thylakoids upon illumination could supply electrons either to PTOX for photoreduction of O_2_ to H_2_O [13] or to PSI through PETC.. To distinguish between these two pathways, we performed CTOE in air-saturated buffer with/without addition of mercuric chloride (HgCl_2_). HgCl_2_ is known to affect electron transfer via plastocyanin (PC) and hinder electron flow from cytochrome b_6_ f (Cytb_6_f) to PSI [21]. PPBQ and FeCy were not added during this measurement as they can take electrons upstream of Cytb_6_f, and hence influence the effect of HgCl_2_. We found that O_2_ consumption in both S and ES was massively diminished (Fig. 2D) compared to without HgCl_2_ supplementation in air-saturated buffer (SI Appendix Fig. S3). Taken together, these results show that O_2_ consumption in the ES samples was a photoreduction process occurring around PSI with electrons supplied from the heavily reduced PQ pool and PTOX was not the major site of O_2_ photoreduction.

### O_2_ consumption in ES thylakoids was accompanied by higher PSI oxidation and higher abundance of flavodiiron proteins

O_2_ consumption around PSI is known to have profound effects on the redox state of PSI [21], [48]. In line with previous reports [29], [34], in our study, P700 became more oxidized (Y[ND] increased) and acceptor side limitation (Y[NA]) decreased in intact ES needles upon increasing illumination compared to S samples (Fig. 3A). Interestingly, when we compared P700 oxidation (P700^+^) from the fast kinetics of the saturating pulse (SP) between spinach leaves and S pine needles (SI Appendix Fig. S6), we found that in S pine needles, P700 did not show a biphasic re-reduction but instead a transient dip. We interpreted this as follows: P700 first reached maximum oxidation upon the SP trigger, then the oxidation state decreased as electrons were supplied from stromal components, but after 100 ms was re-oxidized back to maximum levels. In ES pine, we detected only a very small transient dip in P700 after 100 ms (SI Appendix Fig. S6B) even though more electrons were supplied from the PQ pool (Fig. 2C). This suggests that electrons were taken from the acceptor side of PSI in ES needles, thereby oxidizing P700. Higher P700 oxidation, when the PQ pool is massively reduced (Fig. 2C), could be due to CEF around PSI [34]. However, as pine needles are very thick [49], [50] due their morphology and wax composition, it was not possible to confirm whether the known CEF blocker antimycin A [51] could penetrate the needles. Therefore, to exclude the possibility of CEF mediated P700 oxidation, we monitored the P700 redox state in thylakoid membranes by the SP method in the presence of antimycin A. We found similar results for thylakoids subjected to increasing irradiance, i.e., Y(ND) was higher, and Y(NA) was lower in ES than S in presence of Antimycin A (Fig. 3B).

**Figure 3.**
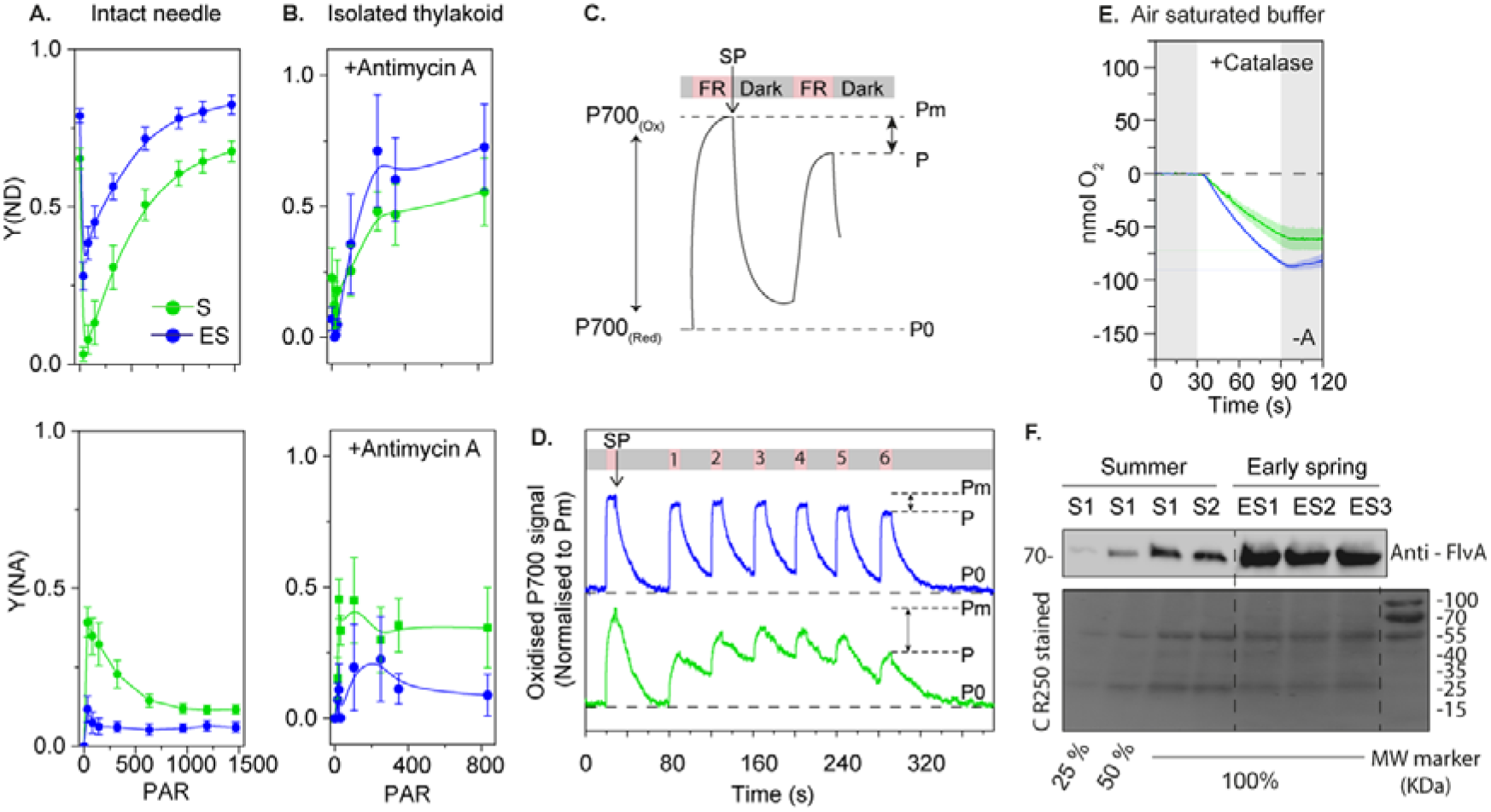
P700 oxidation in S and ES pine thylakoid membranes. **(A)** Changes in donor side limitation [Y(ND)] and acceptor side limitation [Y(NA)] measured by the SP method with increasing PAR (μmol of photons m^−2^ s^−1^) in pine needles (without antimycin A), and **(B**) isolated thylakoid membranes (100 μg/ml) supplemented with 30 μM antimycin A. **(C)** Schematic of SP-induced maximum P700 oxidation (Pm) followed by steady-state P700 oxidation (P) achieved by intermittent FR illumination. The P700 oxidation state was monitored via P700 absorption changes using a DUAL PAM 100 system, where the effective P700 oxidation state was considered as the differences between the Pm and P signal at the 6^th^ cycle of FR illumination. Error bars in A, B indicate the mean (*n*=3) +/− SE. **(D)** Changes in P700 absorbance in S and ES pine thylakoid membranes (corresponding to 100 μg/ml chlorophyll) measured with 6 cycles of intermittent FR illumination in the presence of 30 μM antimycin A (*n*= 3). **(E)** O_2_ yields in S and ES thylakoid membranes in air-saturated buffer measured by a CTOE with 1000 u ml^−1^ catalase supplementation in the absence of ‘A’ (‘-A’). The green/blue colored shaded regions around the O_2_ yield curve indicates ±SE (*n*=3) in the CTOE measurements. In S and ES pine thylakoid membranes data represents O_2_ exchange corresponding to 50 μg of chlorophyll in the CTOE measurements. (For the experimental design, see SI Appendix Fig. S1 and S2,). **(F)** Relative abundance of flavodiiron A protein in S and ES pine thylakoid membranes. Two summer samples (S1, S2) and three early spring samples (ES1, ES2, ES3) (SI Appendix Table S1) corresponding to 4 μg of chlorophyll (100%) were loaded in separate lanes and 1 μg and 2 μg of chlorophyll (25% and 50%) of S1 were loaded in the left two lanes as quality controls, and the gel was immunoblotted against anti-FlvA antibody. (For the relative quantitation of Flv proteins in S and ES, see SI Appendix Table S4). A Coomassie stained membrane is shown in the bottom panel. Similar immunoblotting results were obtained in three independent experiments.

In actinic light (during the SP method), both photosystems are excited, and hence steady-state P700^+^ may be influenced by electrons contributing from the PSII side. Therefore, to determine steady-state P700^+^, we exposed S and ES thylakoid membranes to six intermittent cycles of far-red (FR) light [52] in the presence of antimycin A (Fig. 3C). This method does not reflect the maximum oxidized P700 in terms of Y(ND) like the SP method but only indicates steady-state P700 oxidation changes with increasing cycles of illumination as a consequence of electron flow within PSI (Fig. 3C). The PSI population in S and ES may be different due to different abundance of PSI protein components [29], [30]. Therefore, we normalized the P700 signal to the maximal P700 oxidation level achieved by the initial SP (Pm) prior to the start of the intermittent FR cycles. We found that at the 6^th^ cycle of FR illumination in S samples, the steady-state P700 signal (P) decreased by ~50% compared to Pm, whereas in ES, it only decreased by ~15% (Fig. 3D). Moreover, the relaxation of the P700 signal during the dark periods (after FR illumination) decreased faster and further in ES than in S samples. The dark relaxation of P700 is mainly caused by electron donation from luminal components. Thus, the results suggested that the reduced PQ pool in ES donated more electrons to P700 than in S. Overall, the findings indicated that in ES, a strong electron sink apart from CEF at the acceptor side of PSI consumes the excess electrons donated to P700 from the reduced PQ pool.

MR scavenge electrons from the acceptor side of PSI consuming O_2_ and producing H_2_O_2_, subsequently metabolized through the action of peroxidase/catalases. In isolated thylakoid membranes, stromal components are lacking, and the MR requires external supply of enzymes that could convert H_2_O_2_ to O_2_ and H_2_O. To quantify MR-driven O_2_ consumption in ES thylakoids we performed CTOE measurements in air saturated buffer (without PSII acceptors PPBQ and FeCy) supplemented with 1100 unit ml^−1^ catalase. O_2_ consumption did not change in S thylakoids, but in ES consumption decreased by 35% (Fig 3 E) compared to no catalase addition (SI Appendix Fig S3). This suggested that that MR could only explain a minor fraction of the O_2_ consumption in ES thylakoid membranes, and that another acceptor at the PSI site accounted for a bigger part.

As the evidence showed that none of the other possible mechanisms (damaged PSII, PTOX or photorespiration) could explain the major fraction of the O_2_ consumption in ES thylakoids, we considered the involvement of Flv proteins as genes coding for Flv proteins are present in conifer genomes [37]. Spruce FlvA and FlvB are most similar to type 3 Flv proteins in *Physcomitrella sp*. and *Chlamydomonas sp*. (SI Appendix Fig. S7B), which are known to associate with PCEF via photoreduction of O_2_ around PSI [37], [52], [53]. Photoreduction of O_2_ at the acceptor side of PSI has recently been demonstrated and suggested to be related to Flv proteins that can scavenge excess electron from PSI and keep P700 in an oxidized state (P700^+^) in conifers [37]. In the present study, we quantified the abundance of Flv proteins in S and ES pine thylakoids. We found that FlvA accumulation was ~3 times higher in ES compared to S (Fig. 3 E, SI Appendix Table S4) when samples were loaded based on equal chlorophyll.

### The phenomenon could be reproduced under controlled conditions

We acclimated smaller pines to sub-zero (−8°C) temperatures in a climate chamber and found stronger O_2_ consumption in their thylakoid membranes compared to control (SI Appendix Fig S7A). In presence of HgCl_2_ most of the O_2_ consumption disappeared (SI Appendix Fig S7B) but addition of catalase did not change the O_2_ uptake in any of the samples (SI Appendix Fig S7C), unlike in needles collected from the field (Fig 3E). P700 oxidation was also higher in thylakoid samples from sub-zero acclimated needles compared to control (SI Appendix Fig S7D) and FlvA accumulated more (SI Appendix Fig S7E) suggested that this O_2_ consumption phenomena in climate chamber acclimated samples was identical to the natural environment. This indicated that even though here conditions were less harsh than in the field, the same mechanism was evoked.

### Flv proteins may mediate consumption of O_2_ in early spring also in Norway spruce

In ES, all evergreen plants in the boreal region (including conifers) face similar challenges to protect their photosynthetic machinery. To elucidate whether photoreduction of O_2_ at PSI is a general winter response in conifers, we further explored the possibility of higher O_2_ consumption and correlation of Flv protein accumulation in Norway spruce thylakoids during ES (SI Appendix Fig. S7, 8, 9). Norway spruce is a more challenging study system than Scots pine regarding both its biochemistry and physiology. In our study, biochemical preparations of Norway spruce varied more and the “winter states” were less stable during and after preparations than those of Scots pine. However, our data showed that FlvA protein again accumulated in Norway spruce thylakoid membranes and amounts were ~60% higher in ES compared to S (SI Appendix Fig. S8A). Because the accumulation was lower than in pine, the magnitude of changes in other parameters was expected to be lower. This was indeed the case when ES thylakoid membranes were compared to those in S (SI Appendix Fig. S9A): the steady-state P700^+^ signal decreased by ~30% after the 6^th^ cycle of FR illumination compared to the first cycle. The pattern of fast SP kinetics of P700 oxidation in ES was similar in pine and spruce intact needles (SI Appendix Fig. S9B). CTOE measurements with spruce samples did not work in our hands as the thylakoids sticked to the membrane hindering gas exchange, but n TR-MIMS in the presence of DCMU, ES spruce samples showed stronger ^16^O_2_ consumption but no change in ^16,18^O_2_ (SI Appendix Fig. S9) compared to S samples. From these results, it was evident that Norway spruce behaved similarly to Scots pine and that Flv proteins were most likely involved in P700 oxidation. However, compared to pine, the magnitude was lower in spruce. This suggests that Flv mediated O_2_ consumption under stressful conditions like in ES is a common phenomenon among certain groups of conifers.

## Discussion

Conifers constitute a large fraction of terrestrial biomass, but in comparison with angiosperms, algae and cyanobacteria, they are extremely difficult to study mainly because genetic tools are not sufficiently developed. Photosynthesis studies in conifers are also challenging as their photosynthetic tissue – the needles – are significantly different in morphology and chemical composition than that in other plants. However, we recently demonstrated that conifer needles in the winter elicit a protection mechanism [29] that involves direct energy transfer from PSII to PSI (so-called spill-over), a mechanism whose existence in angiosperms is still a matter of discussion. This protection mechanism may also be associated with phosphorylation of PsbS and triple-phosphorylation of Lhcb1, which trigger thylakoid destacking [30]. Here, we present evidence that a second protection mechanism not yet reported in vascular plants and only present in lower green organisms, namely Flv protein dependent O_2_ photoreduction, also contributes to winter survival of conifer needles by providing protection for PSI.

Since reverse genetic tools are not available for conifers, we used different PETC inhibitors and measured O_2_ exchange in thylakoids isolated from S and ES needles. Furthermore, we correlated O_2_ exchange with protein levels to understand how conifer needles under some conditions exhibit substantial light-dependent net O_2_ consumption instead of O_2_ production as during normal photosynthesis. Four mechanisms – PR, PTOX-mediated chlororespiration, MR and Flv-dependent PCEF flow – could explain this phenomenon. By using PETC inhibitors, we employed an ‘elimination approach’ to distinguish between these four possible pathways. Among them, PR was easily excluded since we observed O_2_ consumption in isolated thylakoid membranes, whereas PR requires a contribution from stromal components. Secondly, Lodgepole Pine in low temperatures has been shown to possess an O_2_ dependent excess energy dissipation capability that was assigned to PTOX [38]. However, when we add HgCl_2_ to block electron transfer through plastocyanin, O_2_ consumption drop significantly suggesting that the consumption occur around PSI and not directly from the heavily reduced PQ pool via PTOX (Fig 2D). Moreover, PTOX can only access PQH2 in the thylakoid lumen when the lumen pH is lower than the stroma [13], however, the ‘sustained winter quenching’ in conifers occur in absence of ΔpH [54], [55] which indicates that PTOX mediated consumption of O_2_ and thereby oxidation of P700 is less likely in conifer thylakoids in early spring. Finally, MR - also known as water-water cycle (WWC) - can only take a very minor fraction of the total electrons in the PETC when CO_2_ assimilation is restricted [56] (such as in winter in conifers [34]). Earlier studies have shown that the magnitude of the reaction in general can be low and may require addition of external exogenous electron acceptor in isolated thylakoid membranes [57] to catalyze the H_2_O_2_ formed via MR. H_2_O_2_ otherwise would damage the iron-sulfur clusters of PSI and increase acceptor side limitation, contrary to Fig 3A. Moreover, MR was previously shown to not be involved winter sustainability of conifers [38] and in lune with this we show that the contribution of MR to net O_2_ consumption in our early spring thylakoid membranes was relatively small.

Therefore, the only remaining mechanism that could explain our data is Flv-mediated PCEF. We acknowledge the fact that we reach this conclusion by elimination of other possible mechanisms. A critical proof – demonstration of decreased oxygen consumption and photoprotection in a mutant lacking Flv proteins – is still not possible to obtain from a conifer; transformation/regeneration protocols and genome editing tools have still not been developed enough. But the fact that genes coding for Flv proteins are among the very few photosynthesis-related genes that are up-, not down-regulated, in Norway spruce in the winter [58], [59] that is also reflected in the higher abundance of the protein levels (Fig 3E, *SI Appendix Fig 7A*) - is a direct evidence supporting our assumption.

PCEF via Flv proteins are known to oxidize P700 by accepting electrons from PSI in cyanobacteria and algae [23]–[25], [27], [52], [60], [61], and the presence of genes coding for Flv proteins in conifer genomes has been noted by others [37]. The exact site of Flv interaction with PSI remains unclear, but our data suggested that in both examined conifers, FlvA accumulated massively in ES thylakoid membranes (Fig. 3E, SI Appendix Fig. S7A). Flv proteins are believed to be soluble proteins, but in pine and spruce, they were retained in conifer thylakoid preparations. Spruce FlvA and FlvB are most similar to type 3 Flv proteins in *Physcomitrella sp*. and *Chlamydomonas sp*. (SI Appendix Fig. S7B), which are known to associate with PCEF via photoreduction of O_2_ around PSI [37], [52], [53]. Therefore, our data suggest that Flvs in conifers act as an electron sink from PSI, readily consuming electrons from the acceptor side during light stress and rapidly reducing O_2_ to H_2_O (Fig. 1C, E, F). In this way, the acceptor side remains in a sufficiently oxidized state to accept electrons from reduced P700, and P700 upon illumination can readily oxidize and donate electrons to F_X_ (Fig. 3B, SI Appendix Fig. S9A). In addition, consumption of O_2_ would reduce the risk of ROS production under conditions where the capacity for ROS detoxification mechanisms is low by creating a lower oxygenic environment around the photosystems. Thus, Flv proteins may have a dual protective function. In the winter, and under other severe stress conditions, needle gas exchange is very low due to the thick cuticle and closed stomata. Hence, any oxygen produced by H_2_O oxidation would accumulate, leading to increased risk of photooxidative stress. Whether this oxygen consumption could give anaerobic conditions *in vivo* is hard to estimate, but in our CTOE experiments 15 minutes of illumination was enough to consume all oxygen in the chamber.

It has been shown that in conifers, P700 remains in a donor side limited condition. In other words, P700 is in a highly oxidated state (P700+) [29], [30], [34]. However, we found that in ES thylakoid membranes, the quinone pool (both PQ and UQ) remains in a massively reduced state (Fig. 2C, SI Appendix Fig. S5). How can P700 remain in an oxidized state when the PQ pool is predominantly reduced? An influx of electrons from the luminal side are passed to the primary stable electron acceptor (F_X_). However, in the presence of Flv protein activity, electrons could be passed to oxygen, preventing F_A_, F_B_ from becoming over-reduced, which would otherwise promote ROS formation [21], [62], potentially causing damage [63], [64]. Hence, Flvs could contribute to photoprotection in conifers, in particular in ES, by keeping P700 in an oxidized state to avoid irreversible PSI damage (Fig. 3A, B, C).

In addition to PCEF, CEF is known to lower the reduction pressure on PSI by accepting electrons from the PETC. Most conifers lack all plastidial *ndh* genes [65] known to be involved in CEF. We found that P700 remained highly oxidized in ES compared to S (Fig. 3A, B), even when the Prg5/Prgl1 mediated CEF pathway was blocked with antimycin A (Fig. 3D, SI Appendix Fig. S8). Therefore, CEF is unlikely to be a dominant pathway for scavenging electrons and oxidation of P700 in ES thylakoids. However, as Pgr5/Pgrl1 has been reported to be more abundant in ES than S [34], along with higher abundance of ATP in winter acclimated pine needles [66], we speculate that CEF may contribute to ATP production via proton gradient formation [67].

A schematic of possible electron flow in ES and S conifer thylakoid membranes is shown in Fig. 4. Upon illumination of S thylakoid membranes, electrons generated from H_2_O splitting to O_2_ reduce the PQ pool and further pass through CytB_6_f to PC and then to P700. Upon receiving electrons, P700 becomes reduced and then under illumination becomes re-oxidized by donating electrons to the electron acceptors (F_X_ to F_A_ to F_B_), which are then taken up by Fd for the forward reactions. In ES **(A)**, although the H_2_O splitting reaction slows down, the PQ pool remains in a highly reduced state **(B)** (most likely with contribution from ndh-2 homolog mediated stromal reduction), which contentiously donates electrons to P700. Upon illumination **(C)**, P700 readily becomes oxidized by donating electrons to acceptors (F_X_, F_A_ and F_B_) **(D)**. As a result, the acceptors (F_X_, F_A_ and F_B_) become highly reduced as electron demand from the forward reaction is limited due to down-regulation of the CBB **(E)**. However, F_X_, F_A_ and F_B_ remain in an oxidized state as the electrons are taken up by Flv proteins **(F)**. Higher Flv activity results in stronger O_2_ consumption, resulting in net O_2_ consumption and donor side limited PSI.

**Figure 4.**
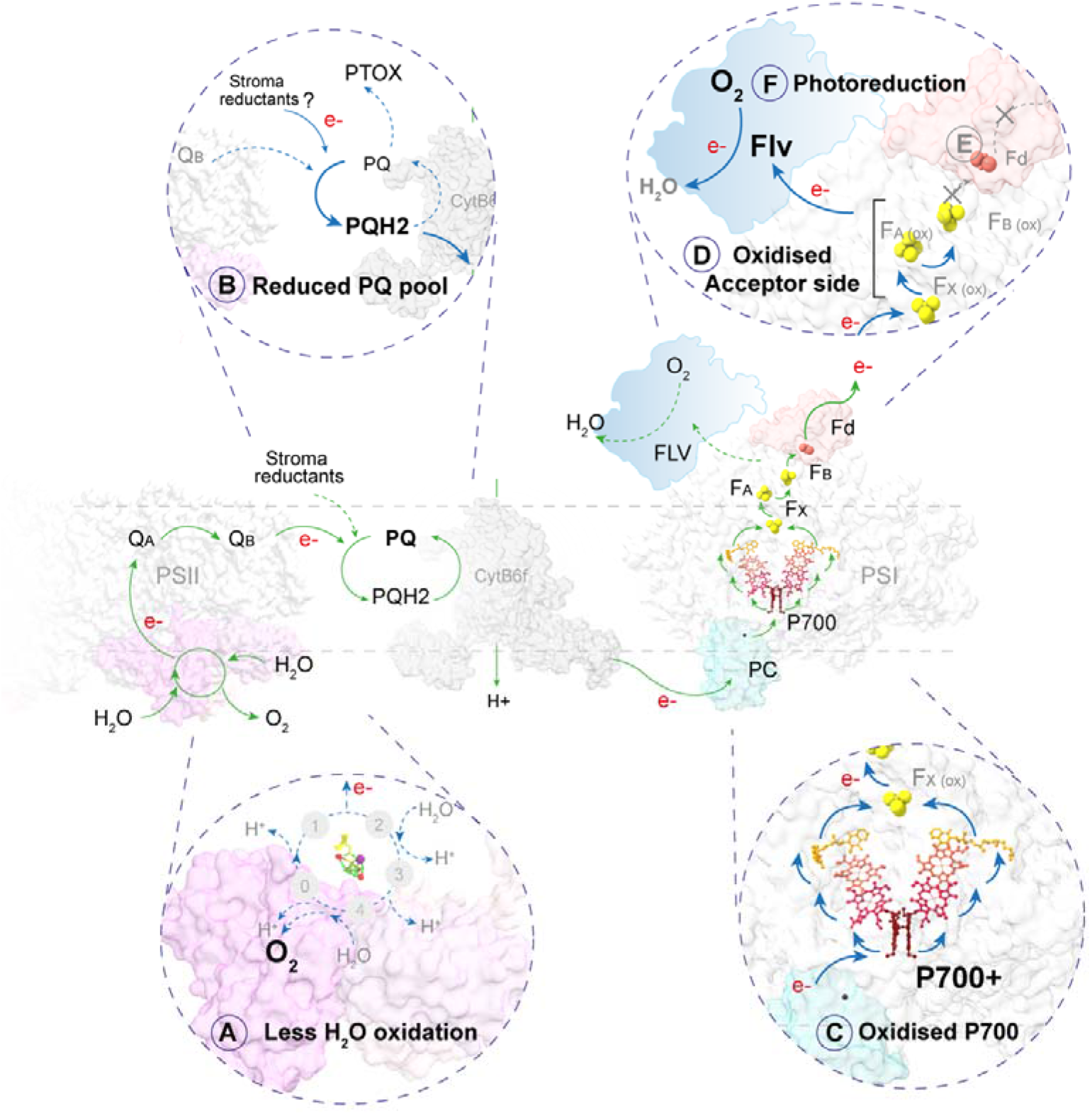
Comparative schematic of light induced electron flow between summer (green) and early spring (blue) thylakoid membranes. Insets show possible alterations in the electron flow in early spring compared to summer as directly measured by our experimental setups. Each step of altered electron flow in early spring is indicated sequentially from **A** to **F**. PDB structures used in the model are: PSII (*Pisum sp*) 5XNL [83]; CytB_6_f (*Mastigocladus laminosus*) 4H0L [84]; plastocyanin-PSI-ferredoxin (*Pisum sp*) 6YEZ [85]. Note that the PSII cofactors in contrast to PSI cofactors, are not shown in the model as their redox state were not measured experimentally. Electron flow from Cytb6f to PSI is mediated through plastocyanin, that upon binding to Cytb6f accepts electrons and diffuses to PSI followed by interaction with PSI and electron donation to P700. However, due to non-availability of Cytb6f bound plastocyanin PDB structure plastocyanin is only shown in association with PSI. The position of Flv on PSI does not represent any empirical binding site nor define the exact the site of electron transfer from PSI acceptorside to Flv. Bold text in the schematic indicates experimental evidences of the phenomena.

The Flv pathway in cyanobacteria and algae is considered ‘primitive’ and consumes a large fraction of electrons from PETC to reduce O_2_, which may lower the photosynthetic yield [37]. However, it was recently shown that Flvs do not compromise CO_2_ assimilation [68]. Perhaps Flvs are useful for conifers as a mechanism that can be rapidly invoked when conditions get worse and dismantled when they get better. In general, having many parallel mechanisms to protect the precious photosynthetic apparatus would be advantageous, and this seems to be the case for conifers. Although we suggest that Flv-mediated oxygen consumption is substantial in conifer needles in ES, this does not mean other mechanisms for O2-reduction are not involved in parallel. Conifers and angiosperms have evolved from a common ancestor that grew in a suitable but light limited environment [69]. Therefore, why have conifers retained but angiosperms lost the Flv proteins? Under low light conditions, carelessly dissipating the precious reducing power by routing electrons to O_2_ would be an evolutionary disadvantage [31]. However, conifers have instead lost the type II NDH mediated CEF and only retained Pgr5/Pgrl1 [65]. It is possible that these different evolutionary trajectories were adaptive as conifers in general adapted to harsh environments but typically competed less well with angiosperms in richer ecosystems. Perhaps Flv proteins provide better protection, but NDH mediated CEF gives better energy economy under a more favorable environment. Hence, Flvs could be a part of a ‘better safe than sorry’ evolutionary strategy in conifers.

## Methods

### Plant material harvesting

Fully developed needles (10-15 gm) were harvested 15 times from the south facing branches of 5-6 mature (40+ years old, 8–10-meter-tall) trees of *Pinus sylvestris* (Scots pine) and *Picea abies* (Norway spruce) (during 2017-2020 (Sampling dates are provided in SI Appendix. Table S1). Needles were immediately transferred to the lab (≤ 5 min) and either subjected to intact needle measurements or thylakoid isolation as described in [35], [70] with slight modifications. After isolation thylakoid membranes were suspended in B3 buffer (50 mM HEPES-KOH (pH 7.5), 5 mM MgCl_2_, 100 mM sorbitol) and either used freshly or flash frozen in 40-50 μl aliquots by liquid N_2_ and stored in −80°C until further use. Before use chlorophyll concentration of each aliquot was determined following [71].

For climate chamber experiments, 15 small 2-3 years old pine trees (1 meter tall) were acclimated to control condition with 120 μmol of photons m^−2^ s^−1^ at 18°C for two weeks and needles were harvested from 5 trees and subjected thylakoid isolation. Then 7-8 trees were moved subzero temperature (−8°C) and acclimated for 10-12 days and needles were harvested and subjected to thylakoid isolation.

### Clark-type electrode measurements

Light induced O_2_ evolution/consumption was measured in a Clark-type electrode (Hansatech instruments, England) with 800 μmol of photons m^−2^ s^−1^ white LED light for 60 seconds (see SI Appendix. Fig S1 for more details) (n=3-4). First background signal was determined by only adding 500 μl of B3 buffer in the Clarke electrode chamber. Later, for all Clarke electrode measurements thylakoid membranes were resuspended (Thawed on ice in case of frozen samples for 30 mins) in B3 buffer supplemented with/without 250 μM PPBQ (2-phenyl-*p*-benzoquinone) and 500 μM FeCy (potassium ferricyanide) as exogenous electron acceptors for PSII [72] to a final volume of 500 μl and measurements were performed immediately (as described in SI Appendix Fig S1A). Temperature was maintained 15°C during all the measurements. For blocking PSII activity, DCMU (3-(3,4-dichlorophenyl)-1,1-dimethylurea) was added at a final concentration of 25 μM from a 100 μM stock solution prepared in DMSO. For blocking electron transfer through plastocyanin, mercuric chloride (HgCl_2_) was added at a final concentration of 1 mg ml^−1^. For catalase activity catalase (Sigma) was added at a final concentration of 1000 unit ml-1 from a stock concentration of 5000 unit ml^−1^ in water. In case of measurements in O_2_-free buffer (N_2_ was bubbled in the CTOE chamber), first 500 μl of B3 buffer supplemented with 250 μM PPBQ and 500 μM FeCy was added in the Clarke electrode chamber and the chamber was closed. N_2_ was supplied in the chamber though a needle from a pressure-controlled gas tap. Once O_2_ signal reached zero, thylakoid samples were added which caused an overshoot in the O_2_ signal. N_2_ Bubbling was continued for 8-10 seconds further to bring the O_2_ signal back to zero. Illumination was started once O_2_ yield reached Zero (SI Appendix Fig S1B). The data shown in all CTOE measurements represent total O_2_ exchange from thylakoid membranes corresponding to 50 μg of chlorophyll.

### Time-resolved membrane inlet mass spectrometry (TR-MIMS) measurements

Gas exchange was measured in thylakoid suspension (in Buffer B3) by TR-MIMS setup as previously described [73]–[75]. MIMS setup contained an isotope ratio mass spectrometer (Delta V ^Plus^; Thermo Fischer Scientific) connected to an in-house built gas-tight membrane-inlet chamber (200-μL) and a cooling trap. Prior measurements, H_2_^18^O (97%; Larodan Fine Chemicals AB) was added (final enrichment of 10%). Analysis of O_2_ reactions was performed on the m/z 32 (^16^O2) and m/z 34 (^16,18^O_2_) signals, with Faraday cup amplification of 3×10^8^ and 1×10^11^ (in the present study, the signal m/z 36 (^18^O_2_) was not analyzed due to its’ low amplitude at the employed H_2_^18^O enrichment and its overlap with the ^36^Ar (m/z 36) signal). Illumination was provided by an 800 μmol of photons m^−2^ s^−1^ external white light source. Recording was started immediately after the samples were loaded in the MIMS cell. Light was switched on 120s after the recording started to achieve the equilibrium of the microoxic environment inside the chamber. The final O_2_ exchange curve was obtained after correction of the gas consumptions by the mass spectrophotometer. The correction was performed by fitting the traces recorded in the dark period before and after the illumination period using Origin Pro 2021 for each individual mass signal (see SI Appendix. Fig S2 for further details of the data analysis).

### P700 measurements

P700 measurements were performed either on intact needles and on thylakoid membranes (n=3) with saturating pulse (SP) (n=3) or on thylakoid membranes (n=3) with intermittent cycles FR illumination by Dual PAM 100 (Walz). First needles or thylakoid membranes were dark adapted from 15-20 minutes. For SP measurements on intact needles P700 absorbance was recorded on a FR light background followed by application of a 600 ms SP of 4000 μmol of photons m^2^ −s^−1^ [48]. For SP measurements on thylakoid membranes P700 absorbance was recorded on a FR light background followed by application of a 50 ms SP of 1000 μmol of photons m^2^ −s^−1^. For measurements on needles PAR (photosynthetically active radiation) was increased step by step upto 1600 μmol of photons m^−2^ s^−1^ whereas for measurements on thylakoid PAR was increased step by step upto 800 μmol of photons m^−2^ s^−1^.

For intermittent cycles of FR light measurement on thylakoids, first, maximum P700 oxidation was determined by a 50 ms SP of 1000 μmol of photons m^2^ −s^−1^, followed by 30s of dark interval and then P700 signal was monitored for 6 cycles of 10 s FR illumination (intensity was 250 μmol of photons m^−2^ s^−1^) followed by 30 seconds of dark interval between each FR illumination [52]. In this measurement thylakoid samples were dark incubated in presence of 30μM Antimycin A for blocking Prg5/Prgl1 mediated cyclic electron flow.

### SDS-PAGE and immunoblotting

Thylakoid membrane proteins were separated on 4-20% TGX PAGE gels (Bio-Rad laboratories) by solubilizing the samples (n=3-4) in Laemmli sample buffer [76] supplemented with 100 μM DDT. After SDS-PAGE, proteins were transferred on a nitrocellulose (Merck) membrane by wet transfer at 15 V for 6h at 4°C and blocked with 2% skimmed milk at room temperature for 2 hours. After blocking, membranes were incubated with FlvA antibody (Dilution 1:2000) over night at 4°C and then incubated with secondary anti rabbit antibody for 2 hours at room temperature and developed using ECL reagent (Agrisera). Image was captured in Azure imaging system.

### Prenylquinones in thylakoid membranes of pine

Three aliquots of the isolated thylakoid samples (~50 μg of thylakoid in 10 μl) were kept in dark (ambient state, N=9) and another set of aliquots were treated with high light intensity (800 μmol of photons m^−2^ s^−1^) for 15 min. Three sample aliquots from the high light treatment were pooled before extraction (N=3). Thylakoids were pelleted down by centrifugation (5000 rpm), sample buffer was removed and three glass beads (3 mm diameter) and 100 μl of ice-cold 30:70 (chloroform:methanol) containing labelled internal standards (Tocopherol) were immediately added to the samples. Samples were extracted for 5 min in a multi-vortex and centrifuged at 4°C at 14 000 rpm for 5 min. Supernatant was transferred into an insert in a vial and samples were analysed by liquid chromatography-mass spectrometry (LC-MS) with a method adopted from [46]. Samples were directly injected in the extraction solvent without drying the aliquots to prevent changes in the redox state of quinones. In addition, quality control samples were prepared by combining an aliquot of each sample group and treatment. Based on the QC samples that were run between sample sets, the state of quinone pool did not change during the whole analysis period. Furthermore, an aliquot of QC sample was mixed with high concentration of ascorbic acid (in methanol, 50 mM) to artificially reduce the metabolite pools and to identify redox active quinones in the thylakoid samples.

The LC-MS system consisted of Agilent 1290 Infinity LC and 6546 LC/QTOF mass spectrometer equipped with atmospheric pressure chemical ionisation (APCI) source and DAD (diode array detector). Metabolites (1 μl sample) were separated with Acquity BEH C18 column (Waters, 100 × 2.1 mm, 1.7 μm particle size) combined with a pre-column. Column temperature was set to 60°C, autosampler temperature to 10°C, and flow rate to 0.5 ml/min. Mobile phase (A=water, B=Methanol) composition at the start was 80% B increasing to 100% B in 1 min, maintained at 100% B for 6.5 min followed by a re-equilibration at 80% B for 1.0 min. The mass spectrometer parameters were optimised with α-tocopherol to increase the sensitivity and to detect mainly molecular ions. The samples were run with positive and negative mode. Data were acquired with a scan time of 4 scans/s and with a range of m/z 120-950. The corona current was 4 μA in positive and 16 μA in negative mode and nebulizer was 45 psi in positive and 40 psi in negative mode. Drying gas flow was 6 l/min, gas temperature 300°C and vaporizer temperature 350°C. Fragmentor was set to 120 V and skimmer to 65 V. In addition, MS/MS spectra were acquired with positive and negative APCI mode for QC samples over a range of collision energy 10, 20 and 40 V. Prenylquinones (11 compounds) were identified based on the standard compounds (α-tocopherol) and MS/MS fragmentation patterns (See SI Appendix. Table S3). Data were processed with Mass Hunter Profinder (version B.08.00, Agilent Technologies). Prenyllipid levels were expressed relative to the total metabolite pool and the redox state based on the ratio between the oxidized (quinone) and reduced (quinol) forms.

### Statistical analyses

The effect of season on quinone redox status (ratio of oxidized and reduced forms) were tested with two-way ANOVA (IBM SPSS statistics version 27). The pairwise comparisons (Post Hoc) were performed with Fisher’s Least Significant Difference test (LSD). In all cases, *P*-value <0.05 was considered significant (See Fig 2D, and SI Appendix. Fig. S5).

Statistically significant differences of FlvA accumulation between samples collected on different dates of the of summer and early spring were calculated by one-way ANNOVA (IBM SPSS statistics version 27) (See SI Appendix. Table S5. Pairwise comparisons were performed with Fisher’s Least Significant Difference test (LSD). *P*-value <0.05 was considered significant.

### Bioinformatic analyses of FlvA and FlvB in spruce

FlvA and FlvB protein sequence were generated from the FlvA and FlvB gene model obtained from the Congenie database (https://congenie.org/) by using Expasy translate tool (https://web.expasy.org/translate/) [77] and was aligned with other flavo proteins in other species (See SI Appendix. Table S4) by using Custal Omega multiple sequence alignment tool (https://www.ebi.ac.uk/Tools/msa/clustalo/) [78]. Unrooted phylogenetic tree was constructed by using IQ-TREE 2 [79] from the aligned sequences. Spruce FlvA and FlvB domains were predicted by using NCBI CDD search tool (https://www.ncbi.nlm.nih.gov/Structure/cdd/wrpsb.cgi) [80] and Expasy PROSITE tool (https://prosite.expasy.org/) [81], [82].

## Acknowledgements

This project was supported by SE2B Horizon 2020 under grant agreement no. 675006 (SE2B) to S.J., the Swedish Research council (VR) to S.J. and J.M., and the Kempe Foundation, FORMAS and SSF to S.J. We would like to thank the photosynthetic platform at UPSC and IRMS platform at KBC (Department of Chemistry, Umeå University) for helpful support with the CTOE and MIMS measurements. We are also grateful to Professor Eva-Mari Aro and colleagues who gave useful comments and, after testing and confirmation, provided antibodies recognizing conifer Flv proteins.

## Supplementary tables

**Table S1.** Sampling and experiment details that has been performed on either intact needles or isolated thylakoid membranes from pine and spruce over three seasons. On each date needles were harvested for either intact needle measurements or for isolation of thylakoid from 5-6 individual spruce and pine trees. For simplicity data only from thylakoid samples from 2018 (Shaded rows) are shown in the manuscript. In case of Clarke electrode, TR-MIMS, P700 absorbance and redox state of PQ pool measurements, data from all ES samples are shown as average of early spring, and data from all S samples are shown as average of summer. Only in immunoblots ES and S dates are shown separately as each lane was loaded with each sampling date. The season specific differences between early spring and summer are consistent in all experiments in all three years.

| Date | Sample ID | Sample | Clarke Elec | MIMS | P700 | PQ pool | Immunoblot |
| --- | --- | --- | --- | --- | --- | --- | --- |
| 2017-02-24 |  | Pine, | Yes | - | Yes | - | - |
| 2017-03-06 |  | Pine, Spruce | Yes | Yes |  | - | - |
| 2017-03-09 |  | Pine, Spruce | Yes | Yes | Yes | - | - |
| 2017-03-17 |  | Pine | Yes | - | Yes | - | - |
| 2017-06-03 |  | Pine | Yes | Yes | Yes | - | - |
| 2017-07-27 |  | Pine, Spruce | Yes | Yes | Yes | - | - |
| **2018-02-12** | **ES1** | Pine, Spruce | Yes | Yes | Yes | Yes | Yes |
| **2018-02-25** | **ES2** | Pine | Yes | Yes | Yes | Yes | Yes |
| **2018-03-12** | **ES3** | Pine, Spruce | Yes | Yes | Yes | Yes | Yes |
| **2018-07-15** | **S1** | Pine, Spruce | Yes | Yes | Yes | Yes | Yes |
| **2018-07-24** | **S2** | Pine, Spruce | Yes | Yes | Yes | Yes | Yes |
| 2020-02-26 | ES4 | Pine, Spruce | Yes | Yes | Yes | - | Yes |
| 2020-03-09 | ES5 | Pine, Spruce | Yes | Yes | Yes | - | Yes |
| 2020-06-21 | S3 | Pine | Yes | - | Yes | - | Yes |
| 2020-07-15 | S4 | Pine, Spruce | Yes | Yes | Yes | - | Yes |

**Table S2.**
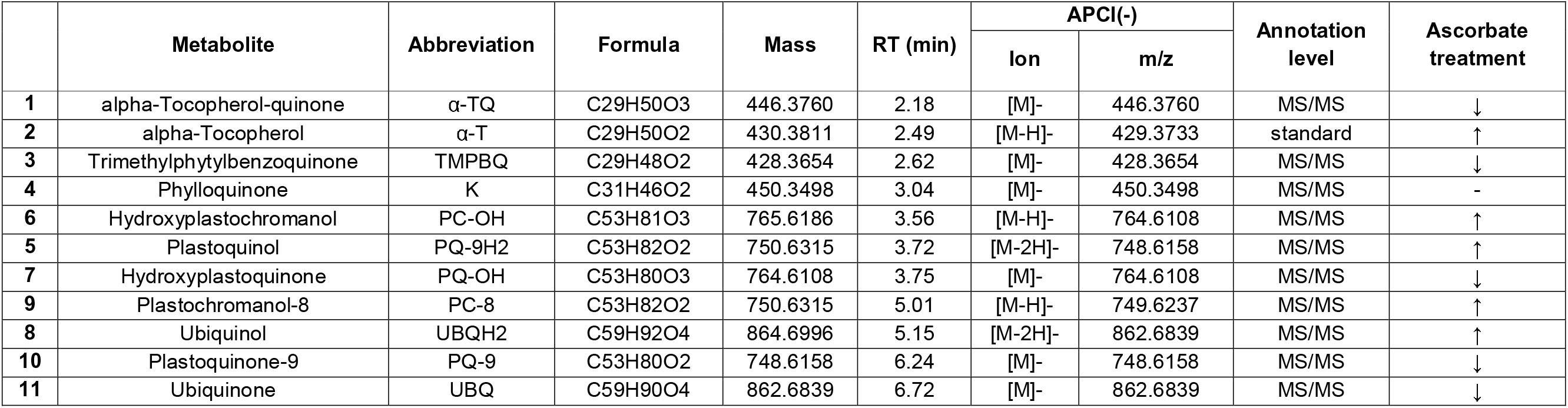
PQ pool metabolite detection. Prenylquinones detected in thylakoids isolated from pine needles using UPLC-APCI(−)QTOF-MS/MS. Ratio of PQH2/PQ are calculated from area under the respective peaks and are shown in Fig. 2D.

**Table S3.**
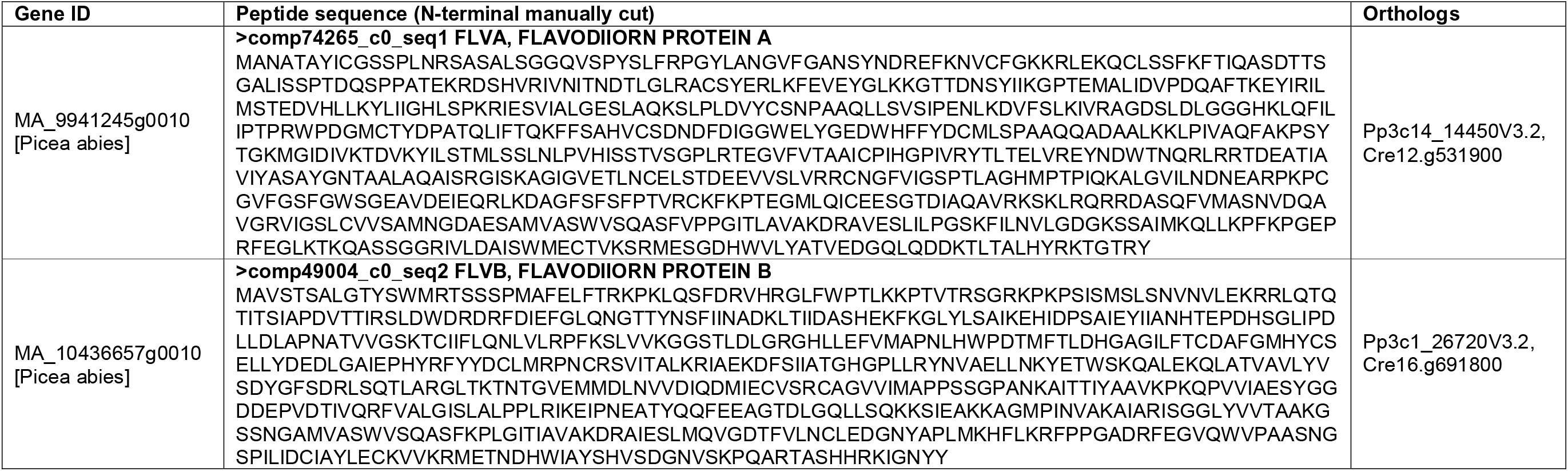
Identity of flavodiiron proteins in spruce. Gene ID, peptide sequence and eukaryotic orthologs (*Physcomitrella patens* and *Chlamydomonas reinhardtii*) of spruce FLVA, FLVB proteins (As shown in the eukaryotic cluster in the unrooted phylogenetic tree of flavodiiron proteins in *SI Appendix, Fig S6B*).

**Table S4.**
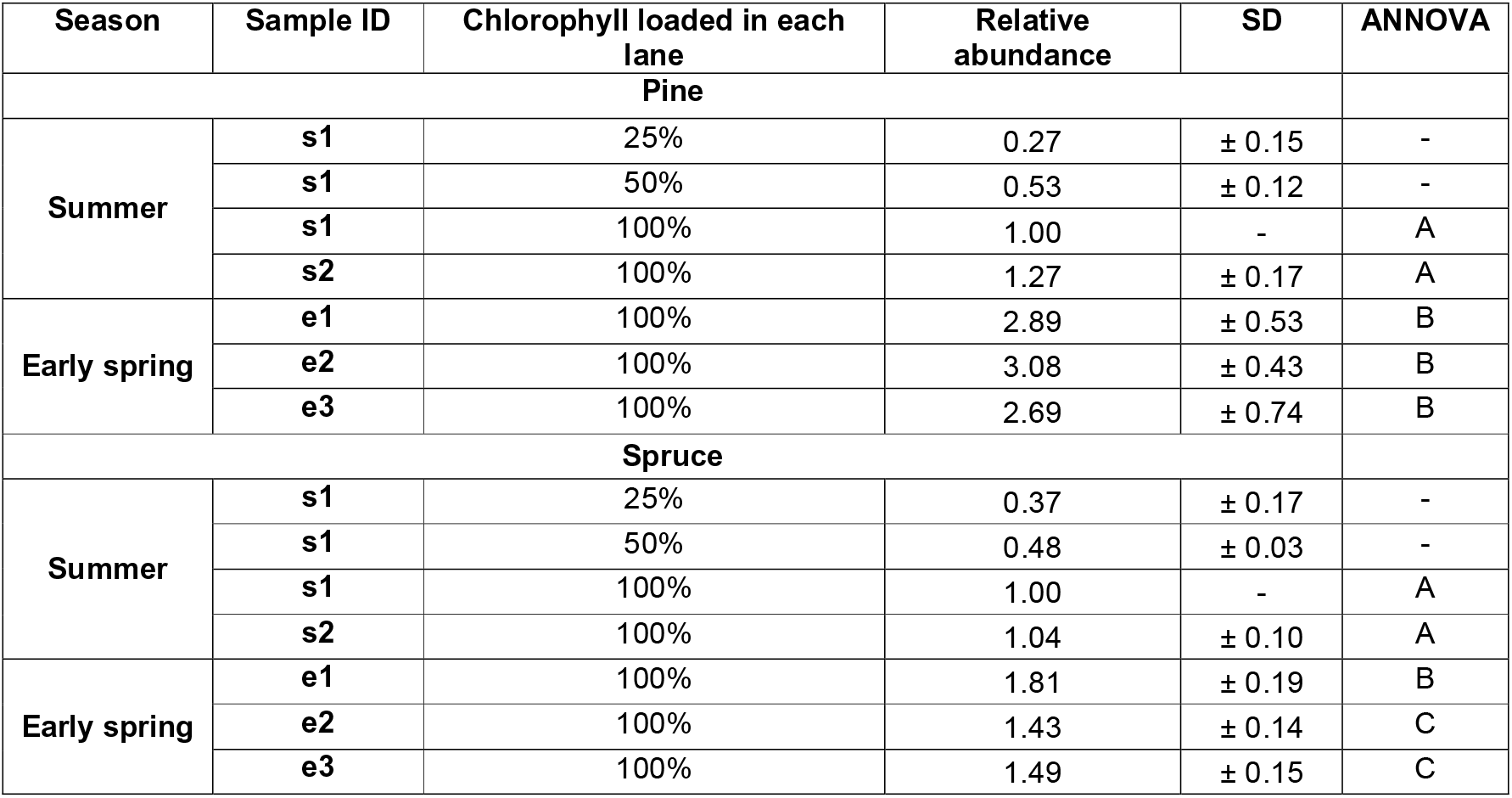
Relative abundance of FlvA protein in ES and S samples in pine and spruce. Here summer sample S1 (2018-07-15) was loaded in 25%, 50% and 100% for quality control, and all other samples loaded on 100% basis, where 100% corresponds to 4μg of chlorophyll in each sample. For quantification, S1 100% band intensity is considered as 1 relative unit (r.u) and rest of the band intensities were normalized based on the 1 r.u. of S1. Avg ± SD is obtained from 4 (For pine) and 3 (For spruce) independent immunoblots. Representative immunoblots of pine and spruce thylakoid are shown in Fig 3D and in *SI appendix, S6A*. Statistical significance was calculated by one-way ANNOVA. Pairwise comparisons were performed with Fisher’s Least Significant Difference test (LSD). *P*-value <0.05 was considered significant (<0.05*, <0.001**)

**Table S5.** Statistical results. The effects of season (summer S vs. early spring ES), treatment (high light, HL) and their interaction on the relative abundance of prenylquinones in pine needles was tested with two-way ANOVA. Pairwise comparisons were performed with Fisher’s Least Significant Difference test (LSD). *P*-value <0.05 was considered significant (<0.05*, <0.001**).

|  | Season | Treatment | Interaction | Season<br>F-value | Season<br>P-value | Treatment<br>F-value | Treatment<br>P-value | Interaction<br>F-value | Interaction<br>P-value | Fisher's LSD |
| --- | --- | --- | --- | --- | --- | --- | --- | --- | --- | --- |
| **PQ** | ** | ** | ** | 12.5 | 2.1E-03 | 34.5 | 9.0E-06 | 28.6 | 3.1E-05 | S-S+HL, S-ES, S-ES+HL |
| **PQH2** | * | ** | ** | 7.9 | 1.1E-02 | 14.8 | 1.0E-03 | 12.9 | 1.8E-03 | S-S+HL, S-ES, S-ES+HL |
| **PQ/PQH2** | * | ** | ** | 8.0 | 1.0E-02 | 15.1 | 9.1E-04 | 13.2 | 1.7E-03 | S-S+HL, S-ES, S-ES+HL |
| **UBQ** | ** | ** |  | 23.1 | 1.1E-04 | 12.8 | 1.9E-03 | 2.8 | 1.1E-01 | S-S+HL, S-ES, S-ES+HL |
| **UBQH2** | ** | ** |  | 14.0 | 1.3E-03 | 9.6 | 5.8E-03 | 3.7 | 6.8E-02 | S-S+HL, S-ES, S-ES+HL |
| **UBQ/UBQH2** | ** | ** |  | 14.4 | 1.1E-03 | 9.7 | 5.4E-03 | 3.7 | 6.9E-02 | S-S+HL, S-ES, S-ES+HL |

## Supplementary figures

**Figure S1:**
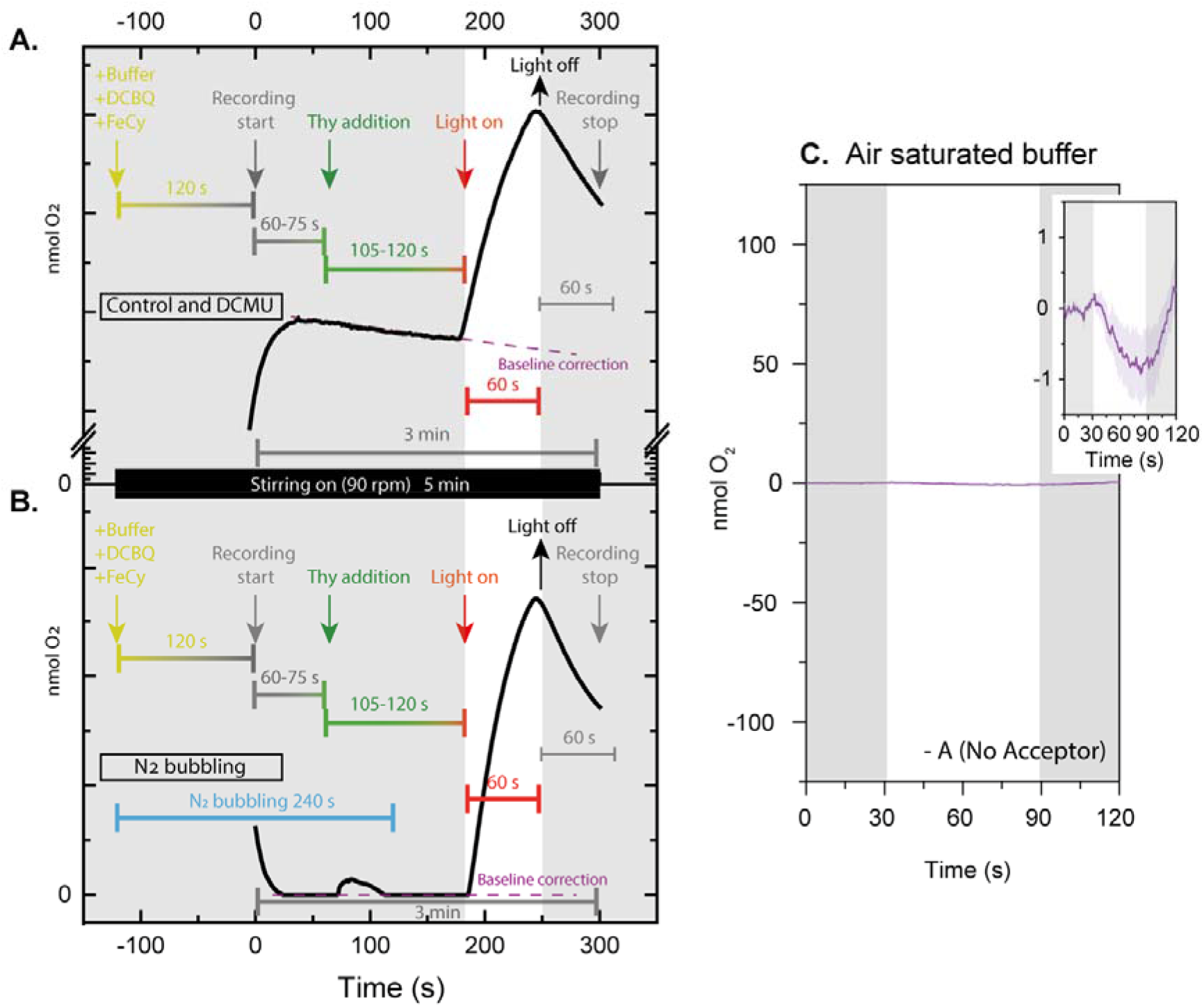
Experimental design for Clark-type electrode measurements on isolated spinach thylakoid membranes suspended in air (A) and N_2_ bubbled (B) buffer in presence (A, B) and absence (C) of exogenous electron acceptor PPBQ and FeCy. Each individual experiment was carried out for 6 mins and 40 seconds. First 250 μl of thylakoid suspension buffer was added in the Clarke electrode chamber, followed by addition of 30 μM PPBQ and 50 μM FeCy. Magnetic stirrer was switched on at a speed of 90 rpm. After 120s instrument recording was started. Thylakoid membranes corresponding to 50 μg of chlorophyll were added after 60s. After 105-120s illumination with 800μmol of photons m^−2^ s^−1^ white LED was switched on. With illumination, increase/decrease in the yield of O_2_ could be observed. After 60s of illumination, LED was switched off and recording was continued for 60s before it was stopped. In case of N_2_ bubbling, after adding PPBQ and FeCy, the chamber was closed and a continuous flow of N_2_ was provided in the Clarke electrode chamber. Thylakoid samples were added after the O_2_ yield reached zero. Addition of thylakoid increased the O_2_ yield slightly, that eventually reached again to zero within few seconds and then N_2_ bubbling was stopped. Then the illumination was started for recording the light dependent O_2_ yields from the thylakoid membranes. Baseline was calculated from the linear range of O_2_ yields prior to illumination but after thylakoid addition to the Clarke electrode chamber. Shaded region following the O_2_ yield curve indicates ±SE (n=3) in CTOE measurements. Inset in **(C)** shows very minor O_2_ activity with 100X zoom in y-axis compared to regular axis in **(C)**.

**Figure S2:**
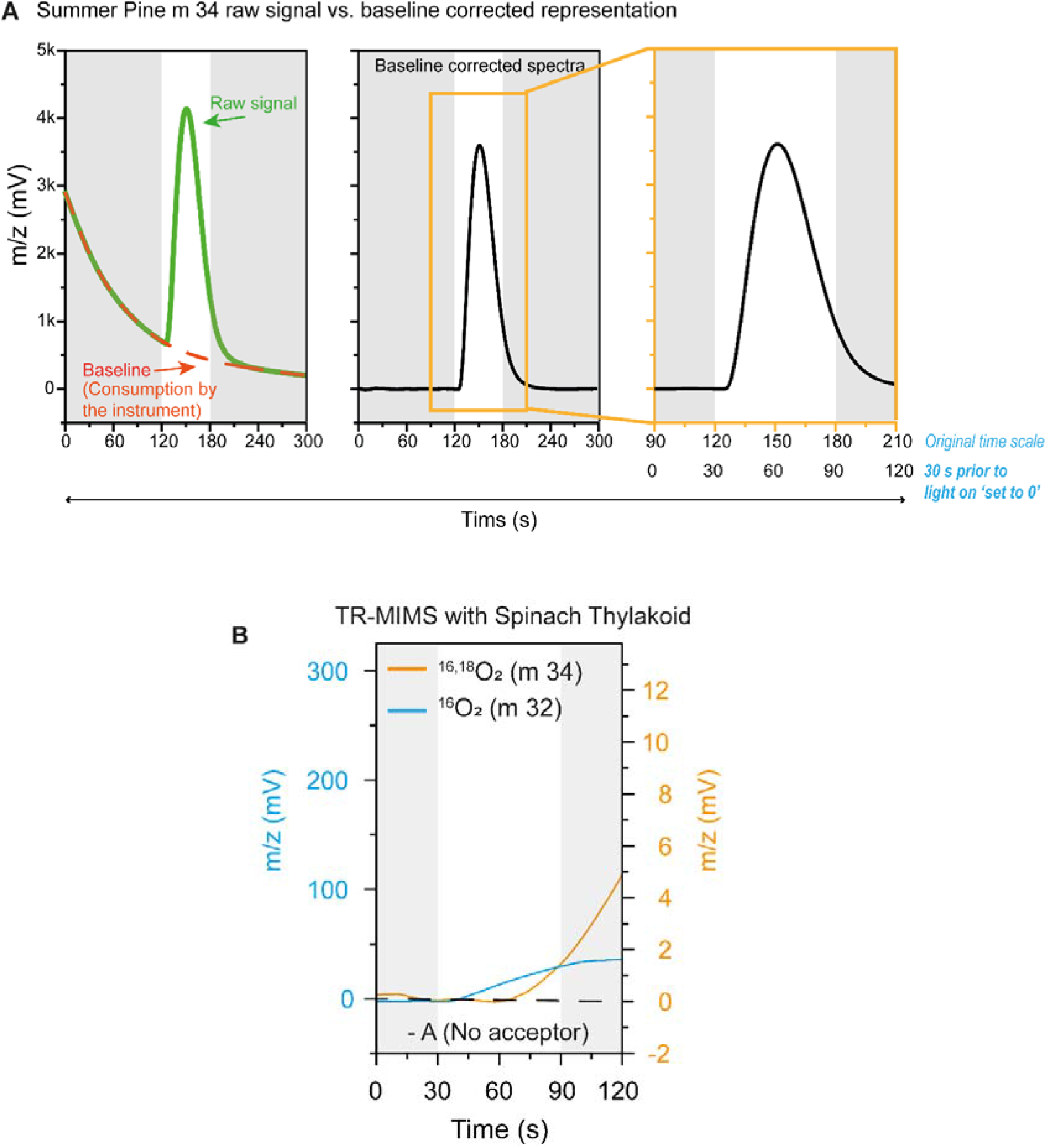
Experimental details of MIMS assays. **(A)** Example of TR-MIMS signal processing. The raw ^16,18^O_2_ signal was recorded for ~ 5 min by using TR-MIMS, where illumination was provided during the ~3^rd^ min (from 120s-180s). The baseline (represents the ^16,18^O_2_ consumption only by the instrument, as there was no mitochondrial fraction detected in the thylakoid samples with immunoblotting against mitochondrial outer membrane protein VDAC1 (voltage-dependent anion channel 1) was calculated using a 2^nd^ polynomial order Savitzky-Golay filter from whole the dark period before illumination and from the last 10s of the whole recording, with a filter window of 3 and threshold was set to 0.05 (In Savitzky-Golay algorithm, each data point is computed from data points within a moving window. If {***f_i_***|,***i*** = **1**, **2**, …, ***N***} be the input data points and if {***g_i_***|,***i*** = **1**, **2**, …, ***N***}, denote the output data points. Each ***gi*** is computed from {***f_m_***|***i*** − ***floor*** (***npts*/2**) < *m* < *i* + *floor*(***npts*/2**). Where ***npts*** is the value of the Points of Window variable. The Savitzky-Golay method performs a polynomial regression to the data points in the moving window. Then ***gi*** will be computed as the value of the polynomial at position i) (Left graph). The baseline was then subtracted from the Raw signal (middle graph) and only 90^th^ s to 210^th^ s was shown in the main and supplementary figures. The scale was set to 0 at the 30s prior to illumination (right graph). The analysis of the data was performed using Origin Pro 2021. **(B)** Effect of no PPBQ addition on ^16,18^O_2_ and ^16^O_2_ yields in spinach thylakoid membranes.

**Figure S3:**
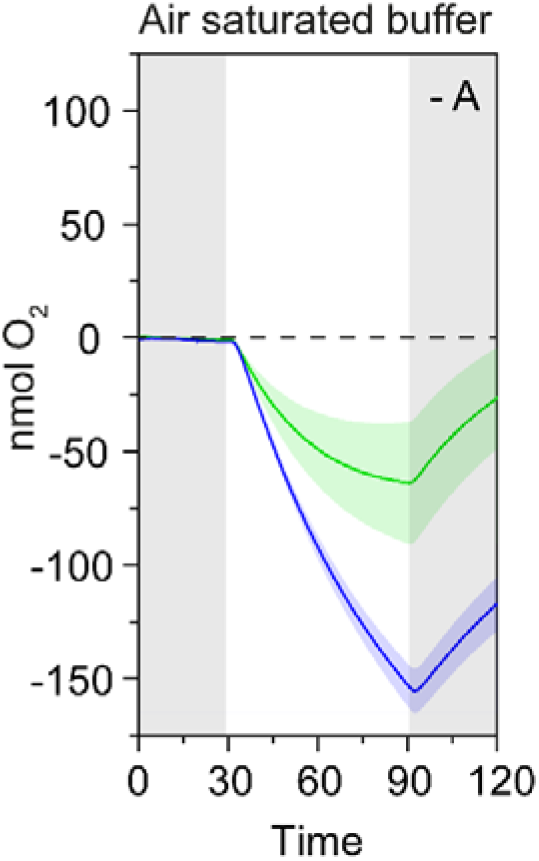
O_2_ yields in summer (S) and early spring (ES) pine thylakoid membranes in air saturated buffer in absence of exogenous electron acceptor PPBQ and FeCy (-A). Shaded region following the O_2_ yield curve indicates ±SE (n=3) in CTOE measurements. In S and ES pine thylakoid membranes data represents O_2_ exchange corresponding to 50 μg of chlorophyll in the CTOE measurements. (For experimental design see SI Appendix. Fig. S1A).

**Figure S4:**
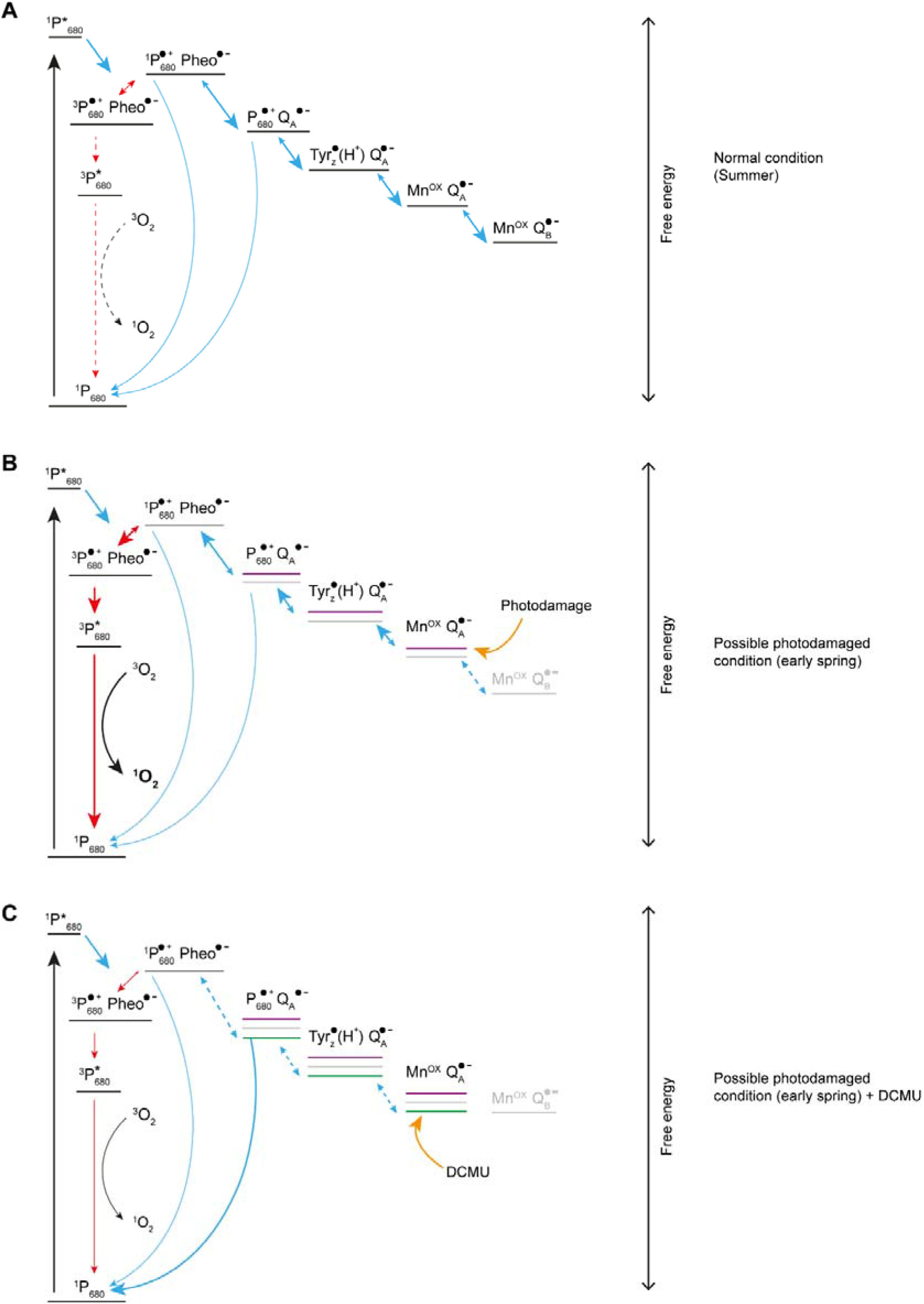
Schematics of electron flow within PSII. **(A)** schematics represent the electron transfer reactions in PSII triggered upon illumination. A series of radical pairs are formed though which electrons are successively transferred forward and ultimately reach Q_B_ site and subsequently reduce a PQ bound to Q_B_ and is released as PQH_2_ in the lumen. Each radical pair has slightly lower energy (on the free energy scale) from the previous one (energy levels are denoted with black lines under each radical pair). Therefore, favors the forward electron transfer. This stabilizes the charge separation within PSII, such as, after excitation of ^1^P_680_ to ^1^P^*^_680_, in step 2, ^1^P_680_^•+^ Pheo^•−^ transfers an electron to P_680_^•+^ Q_A_^•−^ and then then the electron is transferred to the next pair and so on. If transfer of electron from ^1^P_680_^•+^ Pheo^•−^ to next radical pair is limited, then charge recombination may take place and form ^3^P_680_^•+^ Pheo^•−^ (Triplet) via spin conversion. This ^3^P_680_^*^ can convert oxygen to ^1^O_2_ and thereby come back to the ground state, i.e., ^1^P_680_. This triplet formation may also occur due to charge recombination with other electron acceptors, if in any step, the energy gap between any two of the radical pairs decreases then the radical pair may not be able to transfer the electron to the next pair and may transfer electron to the previous radical pair. Therefore, favoring a back reaction and triplet formation and conversion of oxygen to ^1^O_2_,. ^1^O_2_ reacts to proteins leading to damage and this phenomenon leads to O_2_ consumption, which can be measured by monitoring yield of O_2_. **(B)** It has been shown that photodamage of the reaction centre may occur at the donor side (Mn cluster) or at the acceptor side (Q_A_). Under strong white light photodamage majorly occurs at Q_A_, whereas UV light induces Mn cluster damage. Therefore, it can be assumed that natural light induced photodamage mainly occur at Q_A_, as natural sunlight at the earth surface contains much lesser UV. In white light, the redox potential of Q_A_ decreases (Magenta lines) and therefore, in one hand, Q_A_ cannot transfer electron to Q_B_ anymore and on other hand the energy gap between radical pairs become smaller, which in turn favors back reaction and formation of long lived ^3^P_680_^*^. This will favor the chances ^1^O_2_ formation by spin conversion of ^3^O_2_ [64], [86], [87]. **(C)** The redox potential of Q_A_ can be modulated by addition of 3-(3′, 4′-dichlorophenyl)-1, 1-dimethylurea (DCMU), which binds to Q_A_ and increases the redox potential (Green line). This means the energy gap between ^1^P_680_^•+^ Pheo^•−^ to will increase in presence of DCMU. Therefore, the probability of the back reaction form P_680_^•+^ Q_A_^•−^ to ^1^P_680_^•+^ Pheo^•−^ will be very low. Rather the chances of non-radiative charge recombination to the ground states will be much higher. Therefore, production of ^1^O_2_ from oxygen will be lower as well. Similar situation also occurs if the PSII reaction centre is Mn depleted [88]. Therefore, if indeed the ^16^O_2_ consumption in ES samples was a manifestation of damaged reaction centre (Either at Mn cluster or at Q_A_) then, addition of DCMU would decrease the redox potential of Q_A_ and therefore we should see decrease in ^16^O_2_ consumption.

**Figure S5:**
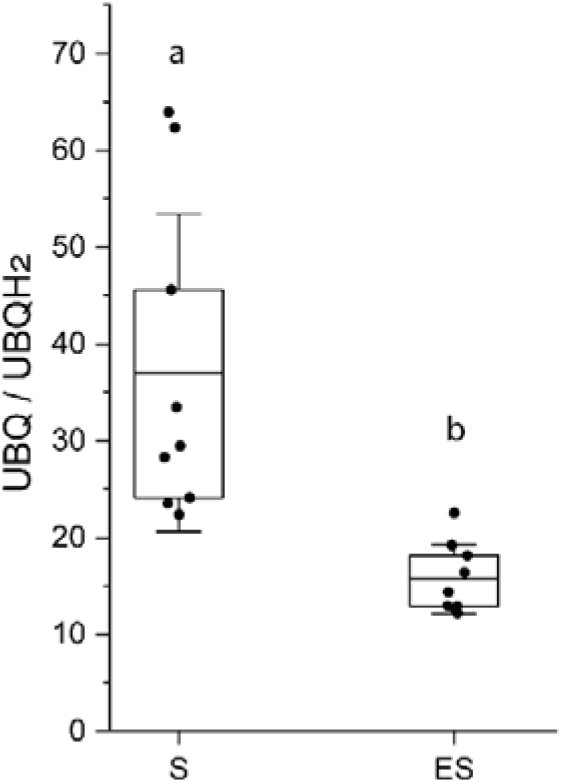
Redox state of the ubiquinone pool presented in terms of ratio between ubiquinol (UQH2) and ubiquinone (UQ) in spruce thylakoid membranes. Ratio of UQ and UQH2 was measured using UPLC-APCI(−)QTOF-MS (n=9) (see SI Appendix. Table S3). Two-way Annova was performed for statistically significant mean differences (see SI Appendix. Table S6 for statistical details).

**Figure S6:**
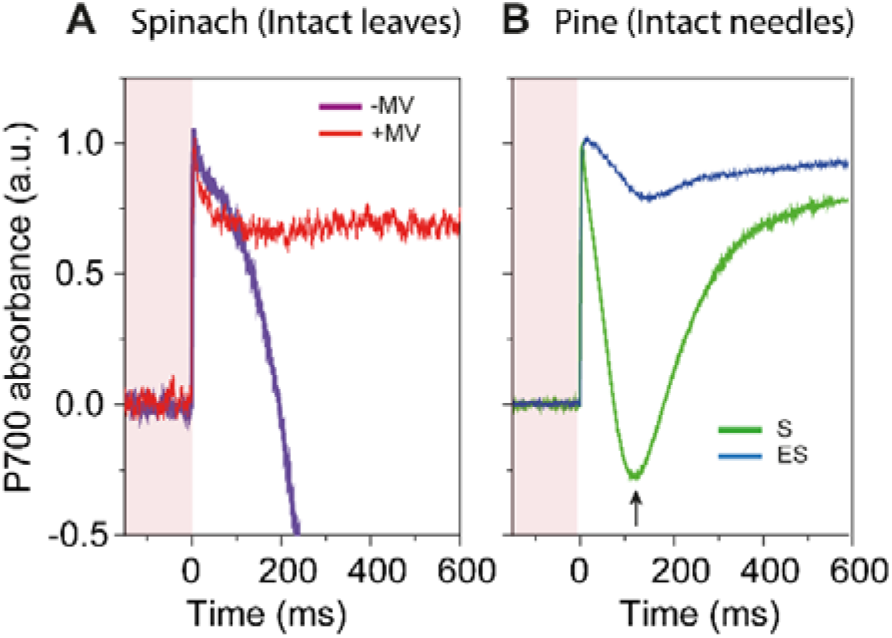
Saturating pulse (SP) induced changes in P700 absorbance in intact pine needles. P700 absorbance was recorded on a FR light background (Red shaded region) with a 600ms SP of 4000 μmol of photons m^2^ − s^−1^ in **(A)** spinach (n=3) and **(B)** pine needles (n=8-12). Spinach leaves and summer pine (S) needles were also vacuum infiltrated with 50μM methyl viologen. (A) In Spinach, upon switching on SP P700 absorbance sharply increases and reached maximum oxidation level within 2-3ms. Then P700 signal dropped for the rest of SP period following a biphasic kinetics because of reduction of P700 via electrons supplied from the luminal components. When MV was vacuum infiltrated prior to measurement, upon switching on SP, P700 dropped only by 15-20% and remain stable for the rest of the SP period. This indicated after maximum oxidation of P700 with SP, even though electrons were supplied from the luminal side but MV being a strong electron acceptor, consumed those electrons rapidly from the acceptor side of PSI and thereby leaving P700 in an oxidized state. **(B)** In S needles (without MV) P700 reached maximum oxidation levels within 2-3ms, and then started dropping as a result incoming of electrons from the luminal side, but approximately after 100ms become re-oxidized (black arrow) and almost reached the maximum level at the end of the 600ms SP period. This typical characteristics of P700 under strong illumination is well characterized as a consequence of electron consumption by flavodiiron proteins in microalgae and cyanobacteria. In early spring needles (ES), the P700 signal behaved almost like MV treated spinach leaves. P700 signal only dropped by 12-15% and then went back to the maximum level. This might be interpreted as lack of electron flow from the luminal side, but as PQ is heavily reduced, therefore, this P700 characteristic strongly indicated that the electrons were provide to P700 from the reduced PQ pool but were immediately taken up by flavodiiron proteins that in turn reduced O_2_ to H_2_O.

**Figure S7:**
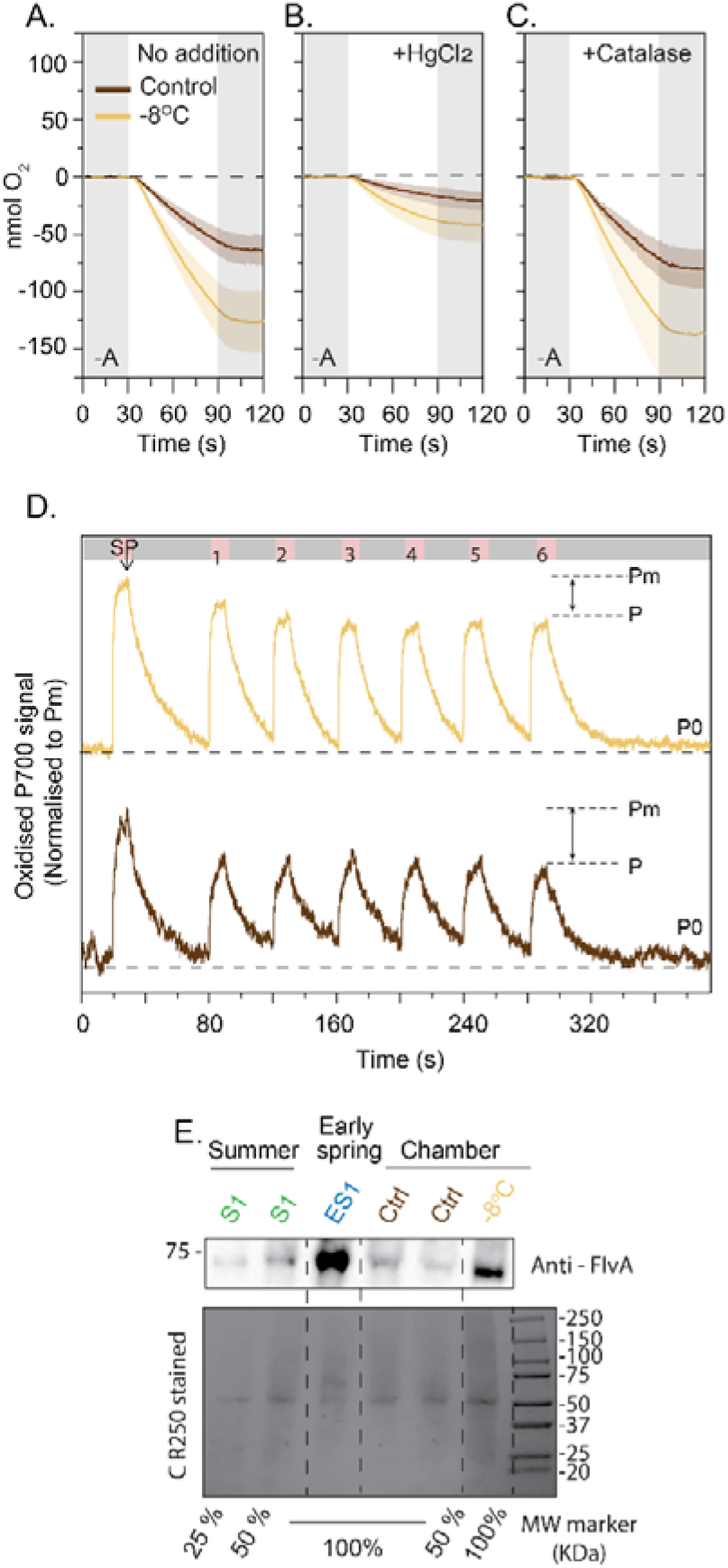
Acclimation of pine tree in low temperature in climate chamber. **(A)** O_2_ yields in Control and sub-zero (−8°C) acclimated pine thylakoid membranes in air saturated. **(B)** O_2_ yields in control and −8°C thylakoid membranes in air-saturated buffer measured by a CTOE with 1 mg ml^−1^ HgCl_2_ supplementation. **(C)** O_2_ yields in control and −8°C thylakoid membranes in air-saturated buffer measured by a CTOE with 1000-unit ml^−1^ catalase supplementation. All CTOE measurements in **(A)**, **(B)** and **(C)** were performed in absence of buffer in absence of exogenous electron acceptor PPBQ and FeCy (+A). Shaded region following the O_2_ yield curve indicates ±SE (n=3) in CTOE measurements. In control and −8°C pine thylakoid membranes data represents O_2_ exchange corresponding to 50 μg of chlorophyll in the CTOE measurements. (D) Changes in P700 absorbance in control and −8°C pine thylakoid membranes (corresponding to 100 μg/ml chlorophyll) measured with 6 cycles of intermittent FR illumination in the presence of 30 μM antimycin A (*n*= 3). (E) Relative abundance of flavodiiron A protein in S, ES, control (Ctrl) and −8°C pine thylakoid membranes. Summer sample S1 and early spring sample ES1, control and −8°C sample corresponding to 4 μg of chlorophyll (100%) were loaded in separate lanes and 1 μg and 2 μg of chlorophyll (25% and 50%) of S1 were loaded in the left two lanes and 50% of control sample was loaded on second lane (from right) as quality controls, and the gel was immunoblotted against anti-FlvA antibody. (For the relative quantitation of Flv proteins in S and ES, see SI Appendix Table S4). A Coomassie stained membrane is shown in the bottom panel. Similar immunoblotting results were obtained in three independent experiments.

**Figure S8:**
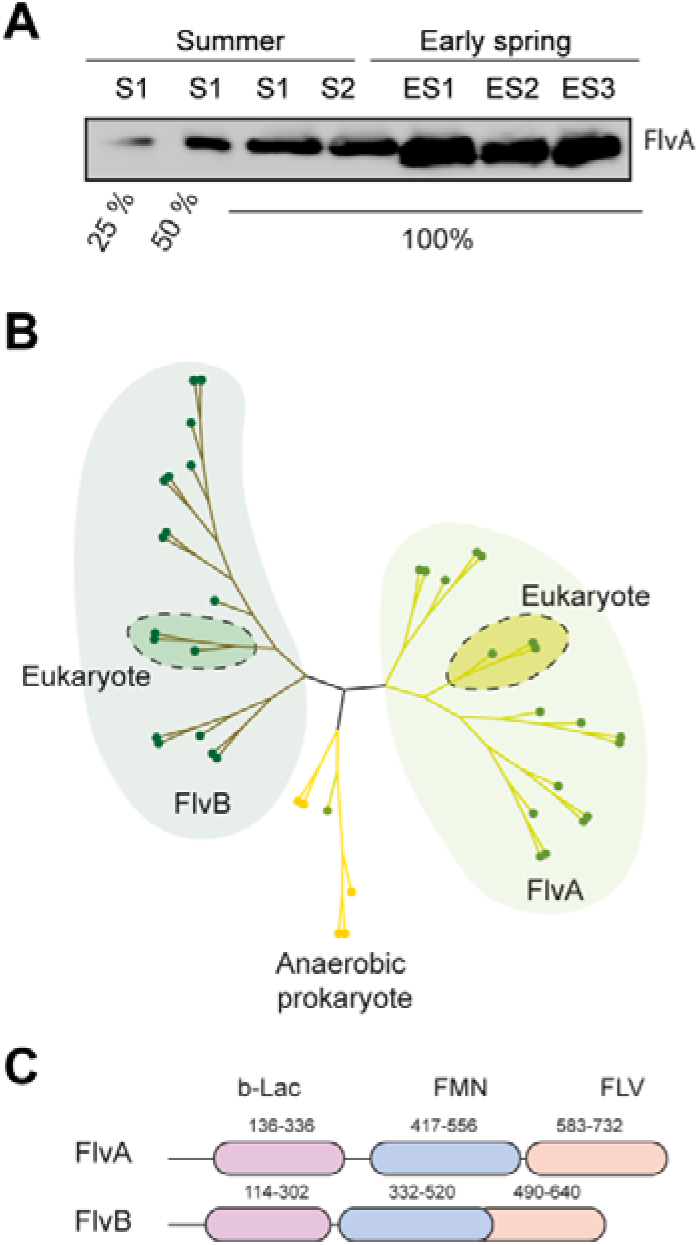
Expression and predicted domains of flavodiiron proteins in spruce. **(A)** Relative abundance of flavodiiron A protein in S and ES thylakoid membranes. Two summer samples (S1, S2) and three early spring samples (ES1, ES2, ES3) (See *SI Appendix. Table. S1* for sample ID details) corresponding to 4 μg of chlorophyll (100%) were loaded in each lane along with 1 μg and 2 μg of chlorophyll (25% and 50%) as quality control in the left two lanes and immunoblotted with Anti-FlvA antibody. For relative quantitation of S and ES see *SI Appendix. Table. S3*. Coomassie stained membrane is shown in the bottom panel. **(B)** Phylogenetic analysis of FlvA and FlvB predicted protein sequences with other flavodiiron proteins from different species (See *SI. Appendix. Table. S4* for list of the Flv proteins in other organisms). **(C)** Domain prediction of spruce FlvA and FlvB proteins. Both FlvA and FlvB contain all three characteristic domains, namely, beta-lactamase domain (b-Lac), flavine mono nucleotide domain (FMN) and flavodiiron domain (FLV).

**Figure S9:**
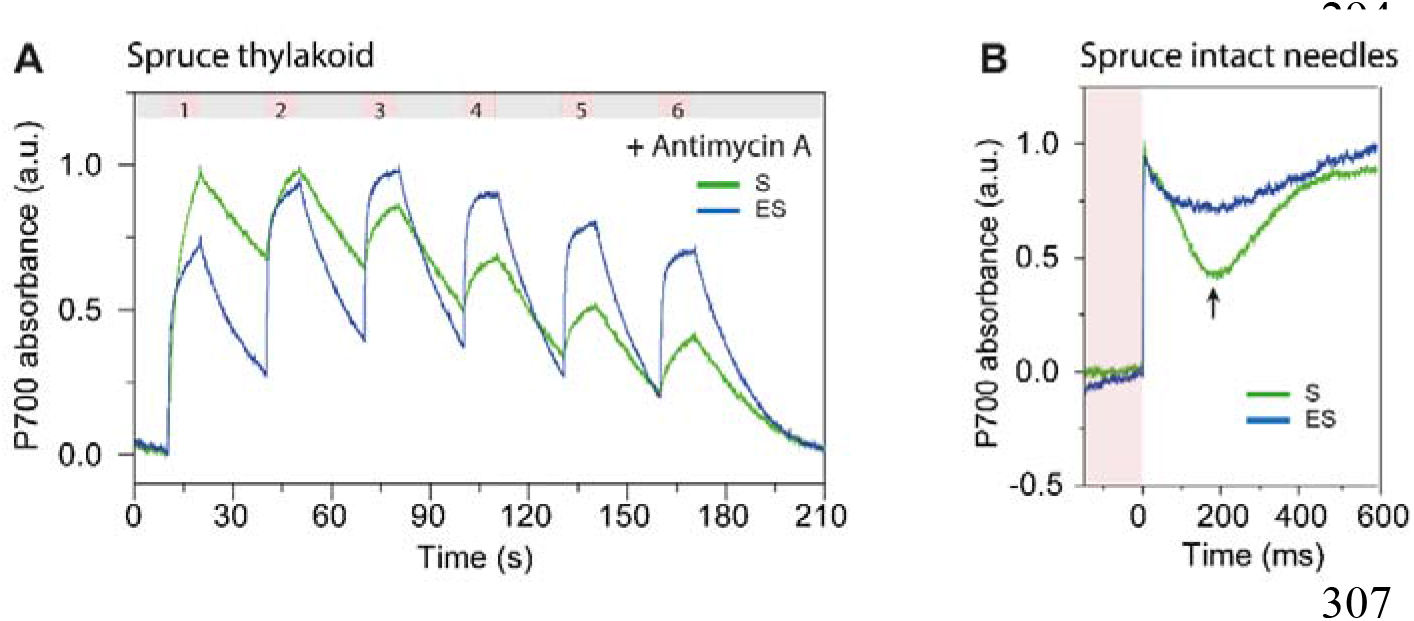
P700 oxidation kinetics in spruce measured with intermittent FR on thylakoid and with saturating pulse (SP) method on intact needles. **(A)** Changes in P700 oxidation in spruce thylakoid was recorded (n=3) by providing consecutive 6 cycles of 10s FR illumination followed by 30s darkness. FR light intensity was 250 μmol of photons m^−2^ s^−1^. Before measurement thylakoid samples were incubated with 30μM Antimycin A, a known cyclic electron flow blocker. **(B)** P700 absorbance changes in intact spruce needles by SP were recorded (n=8-12) as shown in *SI. Appendix. Fig S5* for pine needles.Note that S needles without MV had lower amplitude of P700 reduction after max levels are reached within 2-3ms of SP application. Nevertheless, ES needles behaved similar but not identical to MV treated S needles.

**Figure S10:**
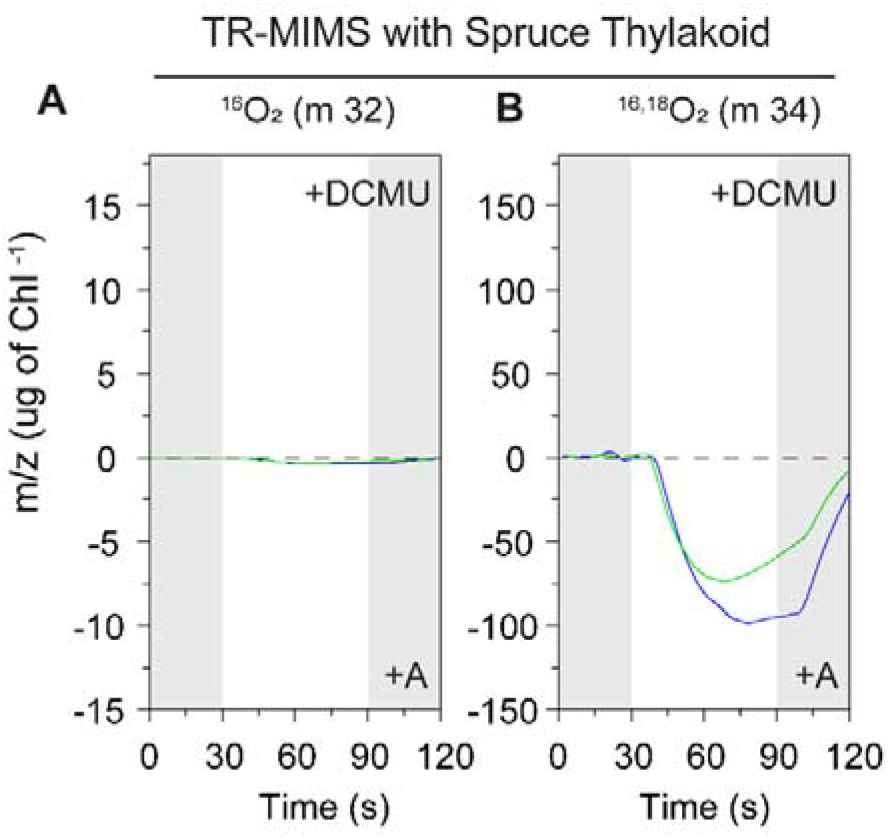
**(A)** ^16^O_2_ and (**B)** ^16,18^O_2_ yields in S and ES spruce thylakoid membranes in air saturated buffer with +A supplemented with 100μM of DCMU measured with TR-MIMS in partially degassed condition. Grey shaded region indicates dark period before and after illumination of the thylakoid membranes with 800 μmol of photons m^−2^ s^−1^ for 60 seconds. S and ES pine thylakoid membranes corresponding to 50 μg of chlorophyll was used in MIMS measurements. (For data analysis details see SI Appendix. S2). For TR-MIMS one representative spectra out of 3-5 independent measurements is shown.

